# Hemodynamic Signals Reshape Biological Inference in Widefield Calcium Imaging

**DOI:** 10.64898/2026.07.03.736453

**Authors:** Tenesha Connor, Macit Emre Lacin, James Hartz, Miguel Malonado, Kemal Ozdemirli, Anthony Sloan, Justin D. Lathia, Sarah Muldoon, Murat Yildirim

## Abstract

Widefield calcium imaging is widely used to study cortex-wide neural dynamics, yet fluorescence signals are strongly influenced by hemodynamic fluctuations arising from blood volume and oxygenation changes. Although hemodynamic correction is frequently applied, it remains unclear whether vascular contributions represent a modest preprocessing concern or systematically bias biological interpretations of cortical activity. Here, we used dual-wavelength imaging to determine how hemodynamic correction reshapes inference of cortical dynamics across mouse lines expressing GCaMP6s, GCaMP6f, and jGCaMP8m, across multiple analytical domains, and under healthy and glioblastoma conditions. We systematically compared uncorrected and corrected signals using analyses spanning functional parcellation, connectivity, spectral structure, brain-behavior coupling, and low-dimensional network-state dynamics. Hemodynamic correction consistently reduced global functional connectivity, increased network modularity, redistributed spectral power away from slow-frequency dominance, and reorganized multivariate representations of cortical state space. These findings demonstrate that vascular signals do not behave as unstructured measurement noise but instead introduce organized variance that propagates across analytical pipelines and influences inference of cortical dynamics. The consequences of this bias were particularly relevant in glioblastoma, where tumor-associated vascular remodeling amplifies the mismatch between fluorescence signals and underlying neuronal activity. In this disease setting, correction revealed hemispheric asymmetries, reduced network-state entropy, and constrained trajectories within cortical state space that were obscured in uncorrected recordings, demonstrating that vascular remodeling can fundamentally alter interpretation of tumor-associated brain dynamics. More broadly, vascular signals systematically biased estimates of functional organization, network architecture, brain-behavior relationships, and disease-associated phenotypes. Together, these findings establish hemodynamic correction as a critical determinant of biological interpretation in mesoscale calcium imaging rather than a simple preprocessing refinement.

## Introduction

Widefield one-photon calcium imaging enables cortex wide measurement of neural population activity in awake, behaving mice, providing a powerful platform for investigating distributed cortical dynamics, functional connectivity, and brain-behavior relationships ^1–7^. By simultaneously capturing activity across much of the dorsal cortex, mesoscale imaging has transformed systems neuroscience by enabling investigation of how large-scale neural networks reorganize during locomotion, arousal transitions, spontaneous behavior, learning, and neurological disease^1,6–19^.

However, fluorescence signals acquired during widefield imaging are not purely neuronal in origin. Hemoglobin absorption and scattering influence both excitation and emission light paths, allowing fluctuations in blood volume and oxygenation to contribute substantially to the measured GCaMP signal ^2,20–24^. These vascular fluctuations are particularly problematic because they are spatially widespread, temporally slow, and strongly coupled to behavioral and arousal-related state transitions ^23,25–30^. Importantly, these signals do not represent unstructured measurement noise, but instead exhibit organized spatial, spectral, and behavioral structure that can bias inferences of cortical activity. Consequently, analyses relying on correlated activity patterns, low-frequency oscillations, or large-scale network organization may be especially vulnerable to hemodynamic contamination embedded within fluorescence measurements.

Despite this limitation, hemodynamic correction practices remain inconsistent across studies, with some pipelines explicitly removing vascular contributions and others omitting correction when calcium dependent fluorescence is assumed to dominate ^5,12,13,31–33^. As a result, it remains unclear whether hemodynamic correction represents a modest preprocessing refinement or a signal-defining step that fundamentally reshapes interpretation of mesoscale cortical dynamics. This uncertainty is particularly important because many commonly used analytical frameworks in widefield imaging, including functional connectivity, spectral decomposition, network modularity, and brain-behavior coupling, rely heavily on slow and spatially correlated activity patterns that overlap strongly with vascular dynamics. Despite the widespread use of these approaches, the extent to which vascular contamination changes biological conclusions across mesoscale imaging workflows remains poorly defined.

This question becomes especially consequential in disease states, where the same vascular physiology that can confound optical signals is itself altered by pathology. Glioblastoma (GBM) is characterized by extensive angiogenesis, abnormal vascular remodeling, blood-brain barrier disruption, and altered neurovascular coupling; all of which profoundly remodel local and global hemodynamics within the tumor microenvironment^34–37^. Additionally, tumor-associated vasculature in GBM is highly heterogeneous and functionally abnormal. These abnormalities alter blood flow, oxygenation, vascular responsiveness, and optical absorption in ways that can strongly influence fluorescence measurements^20,38^. In this context, hemodynamic contamination is not simply a technical artifact superimposed on neural activity but may itself be amplified by disease-associated vascular dysfunction. Consequently, uncorrected widefield imaging signals may incorrectly attribute tumor-driven vascular fluctuations to neuronal dynamics.

Here we evaluate whether hemodynamic correction changes the conclusions drawn from mesoscale calcium imaging in healthy and glioblastoma-bearing mice. Using dual-wavelength imaging and regression-based correction methods, we compare blue-only and hemodynamic-corrected signals across calcium indicators with distinct kinetics and sensitivities (GCaMP6s, GCaMP6f, and jGCaMP8m). We evaluate how correction influences functional organization, connectivity structure, network modularity, spectral composition, brain-behavior coupling, and low-dimensional network-state trajectories. Across analytical domains, hemodynamic correction altered the apparent organization of cortical activity, demonstrating that vascular contributions substantially influence biological conclusions drawn from mesoscale calcium imaging data across both healthy and disease states. Together, these analytical domains encompass many of the principal quantitative frameworks used in contemporary widefield calcium imaging studies, allowing us to evaluate how hemodynamic contamination propagates through an entire mesoscale analysis pipeline rather than influencing isolated metrics.

## Results

### Hemodynamic correction fundamentally reshapes functional organization in widefield calcium imaging

To determine how hemodynamic correction influences both signal properties and inferred functional organization, we implemented parallel regression approaches and evaluated their effects across multiple levels of mesoscale analysis **(Fig. 1; Supplementary Fig. 1).**

**Figure 1.**
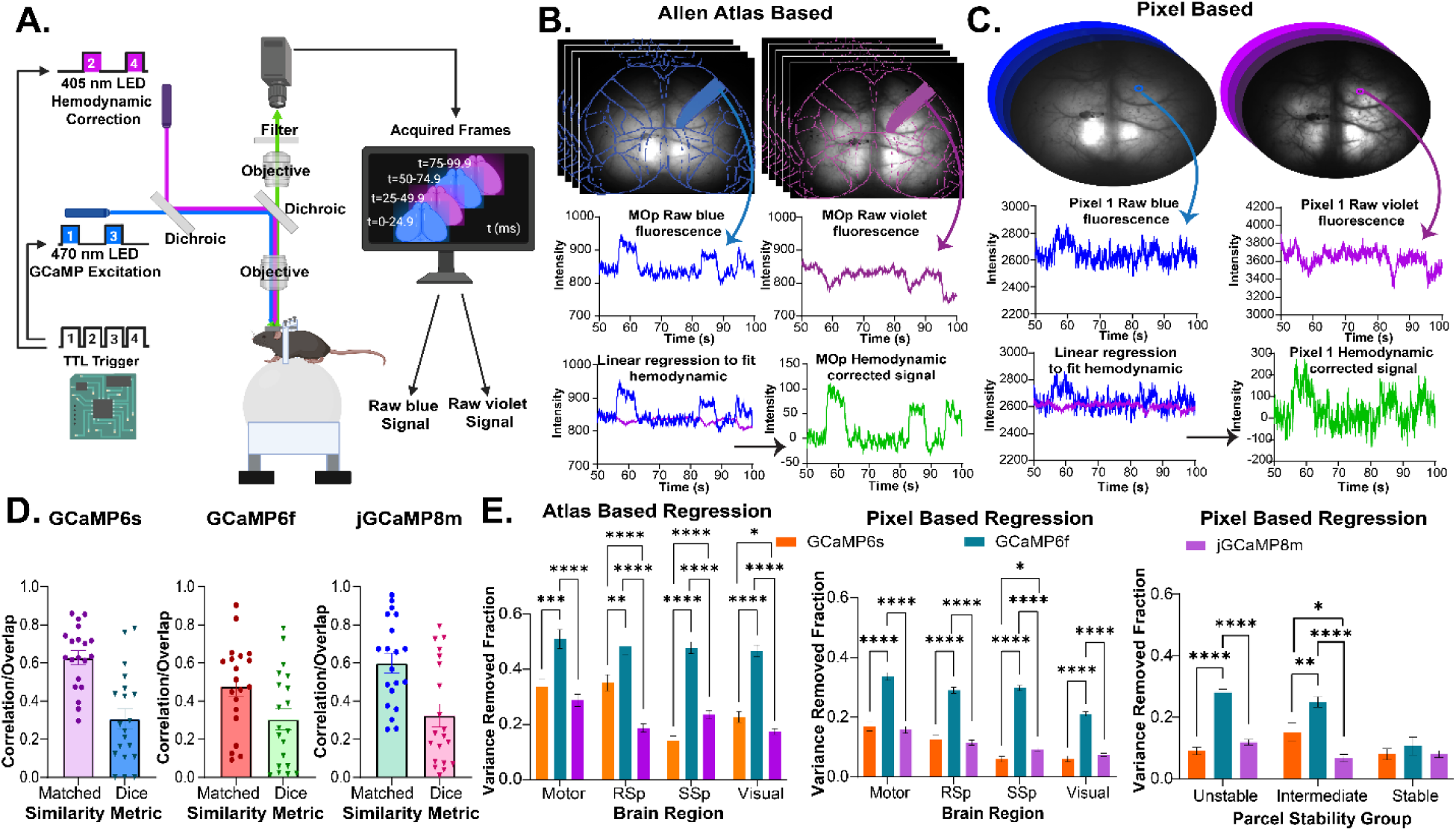
Experimental workflow and analysis pipeline for hemodynamic correction in widefield calcium imaging. **(A)** Dual wavelength imaging system with interleaved 470 nm and 405 nm illumination. **(B)** Atlas-based hemodynamic regression using registered cortical parcels to extract paired blue and violet regional signals. **(C)** Pixel-based hemodynamic regression performed directly on spatially resolved fluorescence signals. **(D)** ICA functional parcellation comparison across GCaMP6s, GCaMP6f, and jGCaMP8m. Each point represents one of 20 ICA derived parcels generated from group level data for each indicator line and evaluated by matched component spatial similarity (correlation) and overlap (dice coefficient), shown on the y-axis. **(E)** Variance removed fraction comparing (left) atlas-based regression, (middle) pixel-based regression grouped by brain region, and (right) pixel-based regression grouped by parcel stability for GCaMP6s (orange), GCaMP6f (teal), and jGCaMP8m (purple). For VRF, replicate sessions were averaged within animal prior to group level summary and bars indicate mean ± SEM across animals. Statistical significance was assessed using multiple paired two-tailed t-test with Holm Šídák correction. Significance is indicated as *p* < 0.05 (*), *p* < 0.01 (**), *p* < 0.001 (***), and *p* < 0.0001 (****).

Dual wavelength widefield imaging was used to acquire interleaved calcium-dependent (470 nm) and hemodynamic reference (405 nm) signals, enabling regression-based separation of neuronal and vascular contributions to the measured fluorescence signal **(Fig. 1A)**. We applied both Allen atlas-based and pixel-based regression strategies **(Fig. 1B-C),** providing complementary approaches for modeling spatially heterogeneous hemodynamic contributions.

We first examined whether hemodynamic correction alters the spatial organization of cortical activity using independent component analysis (ICA)-derived functional parcellations **(Fig. 1D; Supplementary Fig 1A-C).** Comparison of parcellations generated from blue-only and hemodynamic-corrected signals revealed consistent shifts in parcel boundaries and regional segmentation across all GCaMP lines. To quantify these changes, matched ICA components were evaluated using spatial correlation **(Supplementary Fig. 1D-F)** and dice overlap metrics **(Supplementary Fig. 1G-I).**

Across all indicators, spatial correlation values were consistently higher than dice overlap measurements, indicating preservation of coarse large-scale cortical topology despite substantial local reorganization of parcel boundaries. In GCaMP6s mice, matched components exhibited a mean spatial correlation of 0.63 ± 0.04 and dice overlap of 0.31 ± 0.05, indicating moderate structural similarity accompanied by marked boundary reshaping. Similar patterns were observed in GCaMP6f (correlation: 0.48 ± 0.05; dice: 0.30 ± 0.06) and jGCaMP8m (correlation 0.60 ± 0.05; dice: 0.32 ± 0.06). Together, these findings demonstrate that hemodynamic correction preserves large-scale cortical topology while reorganizing local functional boundaries, indicating that vascular signals influence not only measured activity but also functional architecture used to infer cortical organization from mesoscale recordings.

To further characterize the influence of hemodynamic signals on fluorescence activity, we quantified the variance removed fraction (VRF) following regression, where higher VRF values indicate a greater estimated hemodynamic contribution to the recorded signal **(Fig. 1E; Supplementary Figs 1J-R)**.

For atlas-based regression, VRF differed significantly across GCaMP lines and cortical regions **(Fig. 1E (left); Supplementary Fig. 1J-L)**. In general, GCaMP6f exhibited the highest variance removal (mean ± SEM: 0.48 ± 0.01), compared to GCaMP6s (0.26 ± 0.05) and jGCaMP8m (0.22 ± 0.03). Pairwise comparisons revealed significantly greater VRF in GCaMP6f relative to GCaMP6s across all regions (Motor: *p* = 0.0005; RSP: *p* = 0.0026; SSp and VIS: *p* < 0.0001) and jGCaMP8m (*p* < 0.0001 across all regions). Differences between GCaMP6s and jGCaMP8m were more region-dependent, with significant differences in retrosplenial, somatosensory, and visual regions, but not in motor cortex.

Pixel-based regression further revealed consistent differences in hemodynamic contribution across GCaMP lines **(Fig. 1E (middle); Supplementary Fig. 1M-O)**. In this approach, regression was performed at the pixel level, then the resulting signals were subsequently aggregated using the same atlas-defined cortical regions to enable direct comparison with atlas-based regression results. As with atlas-based analysis, GCaMP6f exhibited the highest VRF (0.28 ± 0.03), compared to GCaMP6s (0.10 ± 0.03) and jGCaMP8m (0.11 ± 0.02). These differences were significant across all regions (*p* < 0.0001). In contrast, differences between GCaMP6s and jGCaMP8m were more modest, reaching significance primarily within somatosensory regions.

To determine whether differences in VRF were related to changes in functional organization, parcels were grouped according to their stability, following pixel-based regression. Parcel stability was defined by how closely each corrected component matched its corresponding blue-only parcel, using both spatial correlation and dice overlap. Stable parcels retained similar component structure and spatial overlap after correction, unstable parcels showed poor correspondence, and intermediate parcels showed partial preservation. Stratifying pixel-based VRF by parcel stability revealed a strong relationship between hemodynamic contribution and correction-induced changes in functional organization **(Fig. 1E (right); Supp. Fig. 1P-R).** In unstable parcels, GCaMP6f showed substantially higher VRF than both GCaMP6s (0.28 vs 0.090; adjusted *p* < 0.0001) and jGCaMP8m (0.28 vs 0.12; adjusted *p* < 0.0001), whereas GCaMP6s and jGCaMP8m did not differ significantly. In intermediate parcels, all three indicators separated clearly, with GCaMP6f again showing the highest VRF compared with GCaMP6s (0.25 vs 0.15; adjusted *p* = 0.0072) and jGCaMP8m (0.25 vs. 0.067; adjusted *p* < 0.0001). In contrast, stable parcels showed no significant differences across indicators. Variance removal was greatest in parcels that underwent the largest correction-induced changes in spatial organization, suggesting that hemodynamic signals contribute directly to the functional structure from which cortical parcels are derived.

Together, these results demonstrate that hemodynamic correction not only alters signal variance but also reshapes the inferred spatial organization of cortical activity. The magnitude of these effects depends on both GCaMP indicator properties and the stability of functional parcellation. Because ICA-based parcellation and regional segmentation often serve as foundational inputs for downstream analyses, including connectivity, spectral characterization, and brain-behavior coupling, hemodynamic correction-induced changes in functional organization are likely to propagate across multiple levels of mesoscale network inference and ultimately influence biological interpretation.

### Hemodynamic correction reduces functional connectivity and increases network segregation

To determine how hemodynamic correction alters large scale cortical network organization, functional connectivity was computed from region-averaged fluorescence signals using Fisher z-transformed Pearson correlations and compared between blue-only and hemodynamic corrected conditions **(Fig. 2).**

**Figure 2.**
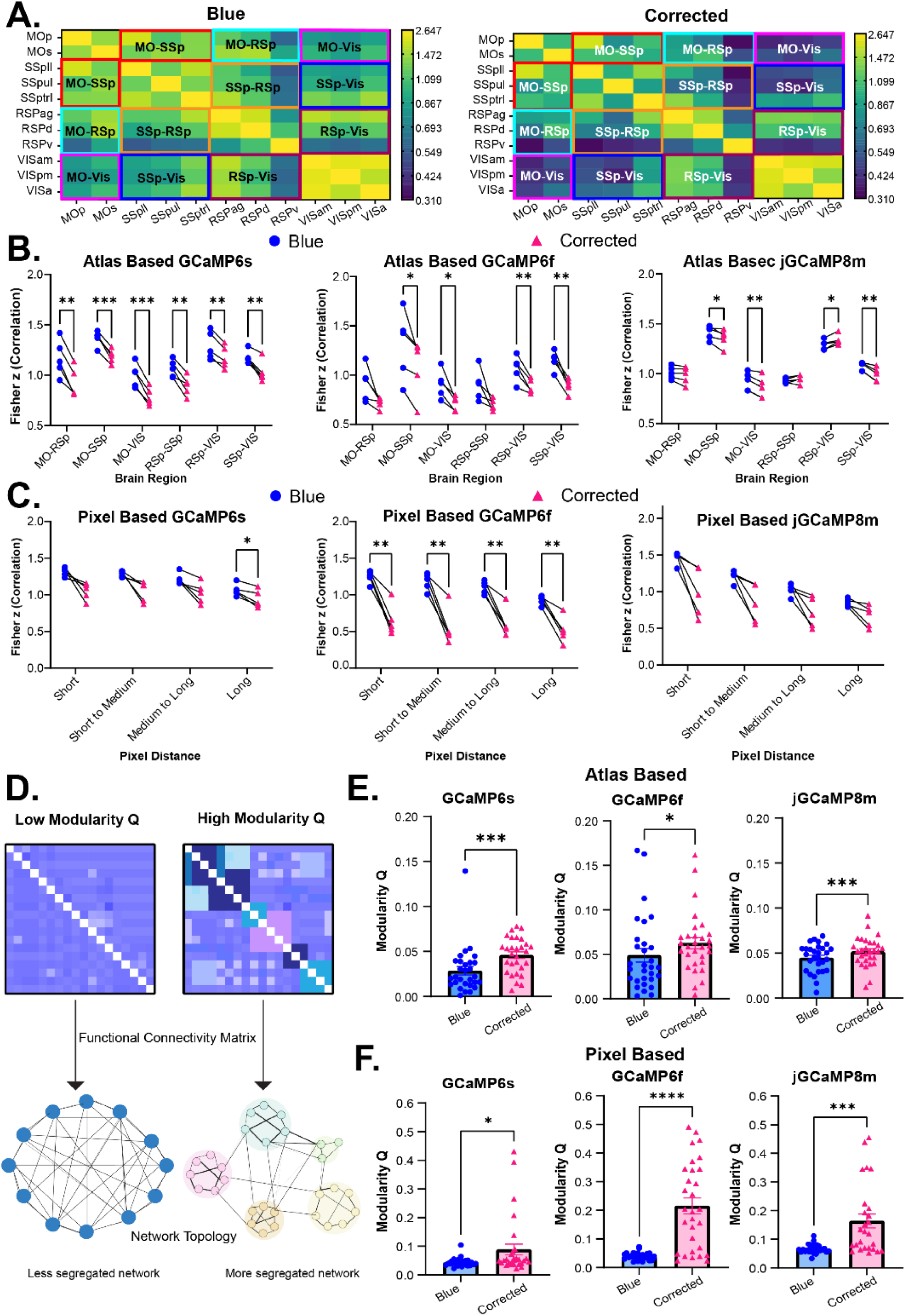
Hemodynamic correction alters functional connectivity and network organization. **(A)** Schematic of region grouping used for connectivity analysis where cortical areas were grouped into motor, retrosplenial, somatosensory, and visual domains. Connectivity was summarized across region pairs for blue-only and hemodynamic-corrected signals. **(B)** Atlas-based functional connectivity (Fisher z) across region pairs for GCaMP6s (left), GCaMP6f (middle), and jGCaMP8m (right). Each point represents a single animal averaged across six sessions. **(C)** Pixel-based functional connectivity of a function of inter-regional distance (short to long range). **(D)** Conceptual illustration of modularity (Q) where functional connectivity matrices are transformed into network representations to quantify community structure and network segregation. **(E)** Modularity (Q) for atlas-based networks across GCaMP lines. **(F)** Modularity Q for pixel-based networks across GCaMP lines. For (E-F), points represent animals from six replicate sessions and bars indicate mean ± SEM. Statistical significance was assessed using multiple paired two-tailed t-test with Holm Šídák correction and paired t-test. Significance is indicated as *p* < 0.05 (*), *p* < 0.01 (**), *p* < 0.001 (***), and *p* < 0.0001 (****).

To facilitate interpretation at the level of functional domains, cortical regions were grouped into motor, retrosplenial, somatosensory, and visual networks and connectivity was summarized across regional pairs **(Fig. 2A).** Across all GCaMP lines, atlas-based functional connectivity was consistently higher in blue-only data compared to hemodynamic corrected signals **(Fig. 2B; Supplementary Fig. 2A-L)** indicating that shared vascular fluctuations systematically inflate estimates of inter-regional synchrony in uncorrected recordings.

In GCaMP6s mice, all region pairs exhibited significant reductions in connectivity following correction (paired t-test, Holm Šídák; *p* = 0.0005-0.0096), with mean connectivity decreasing from 1.18 ± 0.06 to 1.00 ± 0.07 (mean ± SEM). Similar reductions were observed in GCaMP6f, where connectivity was significantly reduced across most regional pairs (*p* = 0.0012-0.0380), with mean connectivity decreasing from 1.04 ±0.07 to 0.83 ± 0.06. In contrast, jGCaMP8m exhibited smaller overall shifts, with fewer region pairs reaching significance and mean connectivity decreasing only modestly from 1.11 ± 0.08 to 1.08 ± 0.09. The smaller correction induced changes in jGCaMP8m suggest reduced overlap between fluorescence activity and the hemodynamic reference signal relative to GCaMP6s and GCaMP6f. Full connectivity matrices confirmed this correction-dependent pattern, revealing widespread reductions in correlation strength across cortical regions after correction **(Supplementary Fig. 2A-L).**

To further examine the spatial organization of these effects, pixel-based connectivity was analyzed as a function of inter-regional distance **(Fig. 2C; Supplementary Fig. 2M-O).** In GCaMP6s, connectivity decreased following correction across all distance scales, although statistical significance was strongest for long-range connections (*p* = 0.0473). GCaMP6f mice exhibited significant reductions across all distance bands (*p* = 0.0085-0.0094), with mean connectivity decreasing from 1.11 ± 0.07 to 0.58 ± 0.03, indicating substantial hemodynamic contributions across both local and long-range spatial scales. In contrast, jGCaMP8m displayed smaller, non-significant reductions despite consistent downward trends across distances. The convergence of reduced functional connectivity and increased modularity after correction indicates that vascular signals contribute broadly distributed shared variance that inflates apparent cortical connectivity, with the strongest effect observed in GCaMP6f recordings.

We next asked whether these pairwise connectivity changes were accompanied by alterations in higher order network organization. Network segregation was quantified using modularity (Q), with higher Q values indicating stronger community structure, defined by dense connectivity within modules and weaker connectivity between modules **(Fig. 2D)**. Across all GCaMP lines, hemodynamic correction significantly increased modularity in Allen atlas-based networks **(Fig. 2E)**, indicating that removal of vascular contributions unmasks more segregated cortical network structure. In GCaMP6s mice, modularity increased from 0.029 ± 0.005 to 0.046 ± 0.004 (*p* = 0.003) following correction, with similar but smaller increases observed in GCaMP6f and jGCaMP8m mice.

Pixel-based networks exhibited larger increases in modularity following correction **(Fig. 2F)**, with significant increases observed across all GCaMP lines (all p ≤ 0.0238) and the strongest effects occurring in GCaMP6f mice. Compared to atlas-based approaches, pixel-level regression revealed substantially greater reorganization of network topology, showing that spatially localized vascular contributions are incompletely captured by Allen atlas-averaged correction strategies.

To visualize edge-level organization underlying these modularity changes, we plotted strong pixel-parcel FC edges relative to corrected-signal community assignments and quantified within- and between-community connectivity **(Supplementary Fig. 3)**. Across GCaMP lines, blue-only networks showed extensive cross-community connectivity, whereas corrected networks showed fewer strong between community edges and a more segregated topology. In GCaMP6s, strong between community edges decreased from 69 in blue-only networks to 27 after correction. Within community edges were more preserved, decreasing from 30 to 26 **(Supplementary Fig. 3A-B)**. This shift was most pronounced in GCaMP6f, where strong between community edges decreased from 109 to 5 and within community edges decreased from 27 to 11 following correction **(Supplementary Fig. 3C-D)**. jGCaMP8m showed a similar pattern, with strong between-community edges decreasing from 77 to 18 and within-community edges decreasing from 33 to 24 after correction **(Supplementary Fig. 3E-F)**.

Overall, corrected networks displayed more clearly defined community structure and reduced inter-module connectivity compared to blue-only data, further supporting the conclusion that hemodynamic contamination obscures modular cortical organization in uncorrected widefield imaging signals.

### Hemodynamic correction removes slow-frequency masking and reveals higher-frequency neural dynamics

Because vascular fluctuations are dominated by slow temporal dynamics, we next examined how hemodynamic correction alters the spectral organization of widefield calcium signals across GCaMP lines (**Fig. 3; Supplementary Fig. 4)**. Frequency-domain analysis revealed that uncorrected signals were heavily biased toward low-frequency activity, whereas hemodynamic correction uncovered higher-frequency neural fluctuations, that were partially masked in the original recordings **(Fig. 3A)**.

**Figure 3.**
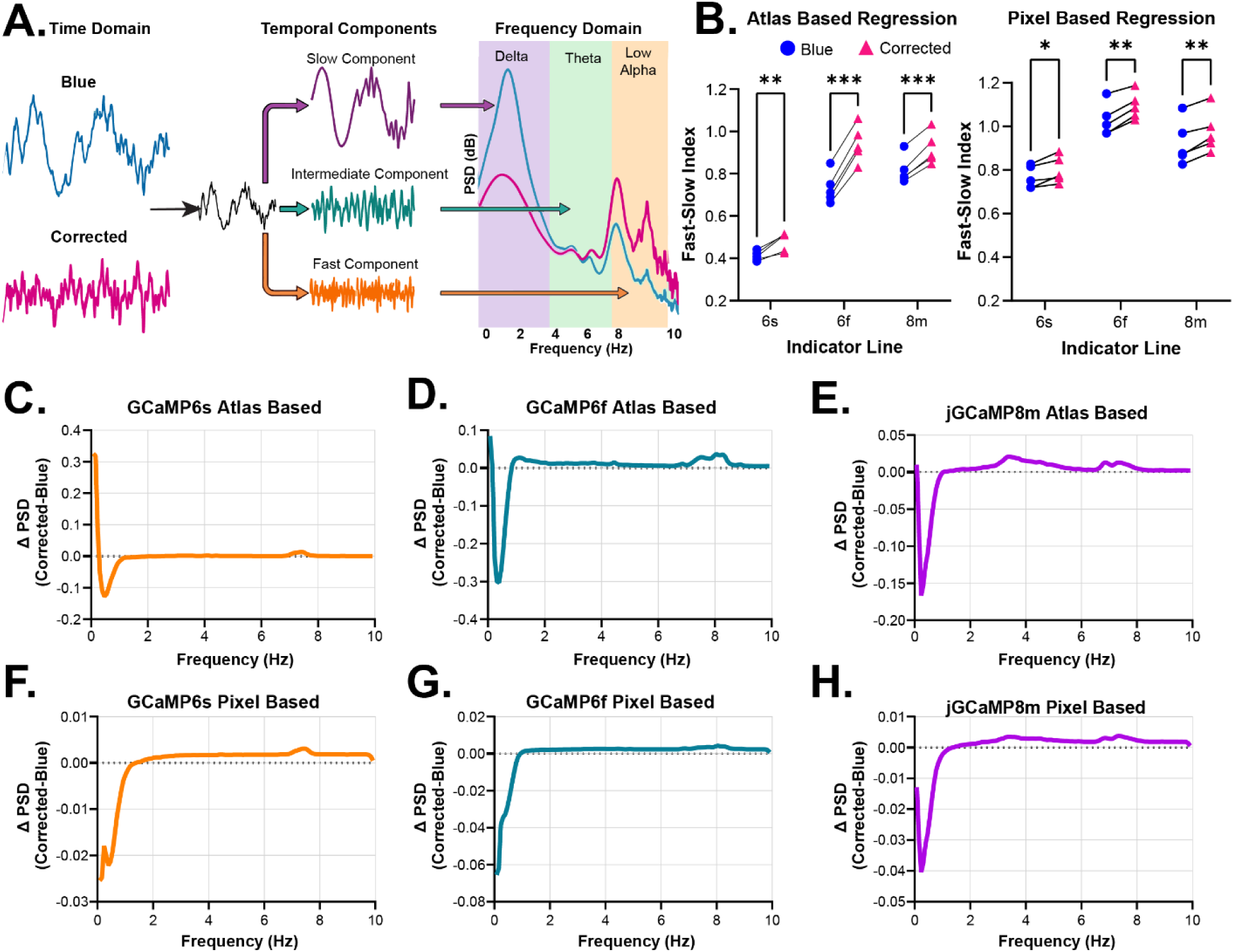
Hemodynamic correction alters frequency-domain dynamics and shifts the balance of slow and fast oscillatory activity. **(A)** Schematic illustrating the transformation from time-domain fluorescence signals to frequency-domain representations. Signals were decomposed into temporal components and analyzed using power spectral density (PSD), with frequency bands defined as delta (0.01-4 Hz), theta (4-8 Hz), and lower alpha (8-9.99 Hz). **(B)** Fast-slow index, defined as *P_θ_* + *P_α_*/*P_δ_*, for atlas-based (left) and pixel-based (right) analyses across GCaMP6s, GCaMP6f, and jGCaMP8m. Hemodynamic correction significantly increased the fast-slow index across all lines (paired two-tailed t-test with Holm Šídák correction; atlas based: *p* = 0.0076, <0.0001, 0.0006; pixel-based: *p* = 0.0218, 0.0039, 0.0060 for 6s, 6f, and 8m respectively). Points represent individual animals averaged across sessions; bars indicate mean ± SEM. **(C-E)** Difference in delta-band PSD (corrected – blue) for atlas-based analyses across **(C)** GCaMP6s, **(D)** GCaMP6f, and **(E)** jGCaMP8m, showing reduced low-frequency power following hemodynamic correction. **(F-H)** Difference in delta-band PSD (corrected – blue) for pixel-based analyses across **(F)** GCaMP6s, **(G)** GCaMP6f, and **(H)** jGCaMP8m, demonstrating similar but more pronounced reductions in low-frequency power at higher spatial resolution. Significance is indicated as *p* < 0.05 (*), *p* < 0.01 (**), *p* < 0.001 (***), and *p* < 0.0001 (****).

To quantify these spectral shifts, we computed a fast-slow index representing the relative balance between higher- and lower-frequency activity **(Fig. 3B)**. Across both atlas-based and pixel-based analyses, hemodynamic correction consistently increased the fast-slow index across all GCaMP lines, indicating reduced dominance of slow (delta) frequency activity following removal of vascular contributions. Specifically, in atlas-based analyses, the fast-slow index increased from 0.73 to 0.94 in GCaMP6f, with smaller but consistent increases observed in GCaMP6s and jGCaMP8m. Pixel-based analyses demonstrated similar trends across indicators, again with the largest shifts occurring in GCaMP6f. The larger correction-induced spectral shift in GCaMP6f shows that this indicator was strongly biased by hemodynamic contamination toward slow-frequency dynamics.

Band specific analyses further demonstrated that these spectral changes were driven by redistribution of oscillatory power across frequency changes **(Supplementary Fig. 4A-F)**. Hemodynamic correction consistently reduced the relative contribution of delta-band activity while increasing theta- and lower alpha-band activity across all GCaMP lines, indicating that vascular signals disproportionately contribute to slow-frequency structure in widefield recordings. These effects were observed in both atlas-based and pixel-based analyses, but were more pronounced at the pixel level, consistent with the increased sensitivity of spatially resolved regression approaches.

Power spectral density (PSD) analyses further illustrated these frequency dependent effects **(Fig. 3C-H; Supplementary Fig. 4G-L)**. Across all indicators, corrected signals exhibited marked reductions in low-frequency power accompanied by relative enhancement of higher-frequency fluctuations. These effects were again most pronounced in GCaMP6f and were consistently observed across both atlas-based and pixel-based analyses. Therefore, hemodynamic correction unmasks faster neural dynamics that are partially obscured in uncorrected widefield recordings by dominant slow vascular fluctuations.

Slow vascular signals are spatially widespread and temporally correlated across cortex, so their removal not only reshapes spectral structure but also alters large-scale functional organization. Hemodynamic contamination therefore biases blue-only widefield mesoscale calcium imaging toward slow-frequency, spatially shared activity, particularly obscuring faster neural dynamics and altering interpretation of oscillatory structure across cortical networks.

### Multivariate analysis reveals coordinated reorganization of brain-state representations following hemodynamic correction

Hemodynamic correction altered connectivity, network modularity, and spectral structure, raising the possibility that these changes reflected coordinated reorganization of cortical activity states rather than independent shifts in individual metrics. To address this, we integrated features across analytical domains within a unified multivariate framework using principal component analysis (PCA) **(Fig. 4; Supplementary Fig. 5)**. Across all GCaMP lines, blue-only and hemodynamic corrected signals occupied distinct regions of low-dimensional feature space, demonstrating that correction reshapes the inferred organization of mesoscale brain states rather than altering connectivity, modularity, or spectral metrics in isolation.

**Figure 4.**
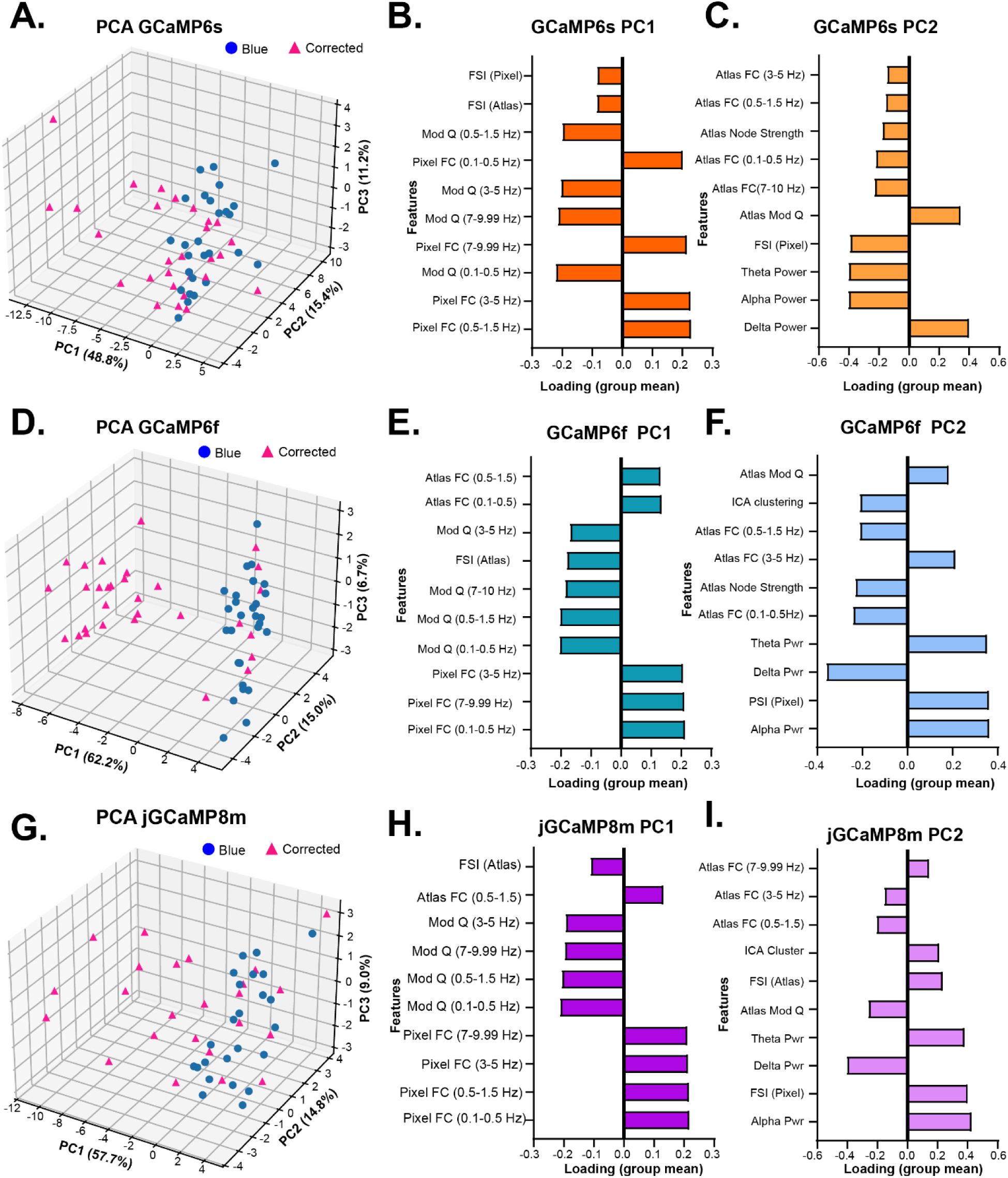
Multivariate analysis reveals coordinated shifts in connectivity, spectral, and network features following hemodynamic correction. **(A, D, G)** Three-dimensional principal component analysis (PCA) representations of session-level feature space for **(A)** GCaMP6s, **(D)** GCaMP6f, and **(G)** jGCaMP8m. Each point represents a single session, with blue only (blue circles) and hemodynamic corrected (pink triangles). The first three principal components explained 48.8%, 15.4%, and 11.2% of variance in GCaMP6s; 62.2%, 15.0%, and 6.7% in GCaMP6f; and 57.7%, 14.8%, and 9.0% in jGCaMP8m. **(B-C)** Feature loadings for **(B)** PC1 and **(C)** PC2 for GCaMP6s, showing the relative contribution of connectivity, spectral, and network features to each axis. **(E-F)** Feature loadings for **(E)** PC1 **(F)** and PC2 for GCaMP6f. **(H-I)** Feature loadings for **(H)** PC1 and **(I)** PC2 for jGCaMP8m. Abbreviations: FC, functional connectivity; FSI, fast-slow index.

In GCaMP6s, session-level representations showed partial overlap between blue-only and corrected conditions but exhibited a consistent directional shift following correction **(Fig. 4A)**. This displacement was primarily aligned with PC1, indicating correction altered a dominant multivariant feature axis rather than producing isolated changes in individual metrics alone. Quantification of Euclidean displacement in PCA space revealed modest but variable shifts across sessions (mean displacement 4.43 ± 2.02, SEM), consistent with a more heterogeneous influence of hemodynamic contamination in this indicator line.

Feature loading analysis demonstrated that the first principal component in GCaMP6s was primarily driven by broadband pixel-level functional connectivity with opposing contributions from network modularity **(Fig. 4B)**. The second principal component captured spectral organization, separating slow (delta) frequency activity from higher-frequency oscillations and the fast-slow index **(Fig. 4C)**. The multivariate analysis in GCaMP6s therefore captured hemodynamic correctional shifts in feature space along axes defined by reduced global connectivity, increased network segregation, and greater relative contribution of faster oscillatory dynamics.

GCaMP6f exhibited the clearest separation between blue-only and corrected sessions in PCA space, indicating a stronger and more consistent effect of hemodynamic correction across animals **(Fig. 4D)**. Euclidean displacement magnitudes were larger than those observed in the other GCaMP lines (mean displacement 7.16 ± 1.48, SEM) and displayed relatively low inter-session variability. Feature loading revealed this separation was driven primarily by broadband pixel-level connectivity **(Fig. 4E)**, whereas spectral features were comparatively weaker **(Fig. 4F)**.

JGCaMP8m also demonstrated a consistent directional shift following correction **(Fig. 4G)**, with intermediate displacement magnitudes of 5.17 ± 1.71 (SEM), indicating stable reorganization of the multivariate feature space across sessions. Similar to GCaMP6f, the leading component was dominated by broadband connectivity structure **(Fig. 4H)**, whereas the second component reflected combined contributions from spectral and network-level features **(Fig. 4I)**, indicating a more distributed feature organization compared to GCaMP6f.

Across all indicators, feature loadings consistently demonstrated the primary axes of variance were jointly defined by functional connectivity, network modularity, and spectral power. Higher order organization was similarly preserved across lines, with the third principal component primarily reflecting atlas-level network integration and organization **(Supplementary Fig. 5A-C)**.

A combined PCA across all GCaMP lines further demonstrated that baseline analysis states differ systematically between indicators, with each line occupying distinct regions of the shared feature space **(Supplementary Fig. 5D)**. These separations reflected intrinsic differences in connectivity structure and spectral composition across GCaMP lines. Despite these indicator-specific starting points, hemodynamic correction shifted all lines in a consistent direction, indicating that correction acts on a shared axes of mesoscale organization dominated by broadband connectivity and large-scale network structure.

Multivariate integration showed that correction-dependent changes were coordinated across analytical domains rather than confined to a single metric class. Blue-only and hemodynamic corrected data separated in a shared feature space, with correction induced shifts aligned along axes weighted by broadband connectivity, modularity, and spectral balance. Thus, hemodynamic contamination was expressed as structured variance spanning multiple downstream measures, changing the inferred organization of mesoscale brain states.

### Hemodynamic correction strengthens brain-behavior coupling while preserving temporal structure

Pupil diameter provides a robust indicator of arousal-linked brain states while pupil-linked widefield fluorescence signals can reflect a mixture of neural activity, vascular dynamics, and autonomic physiology. This makes pupil-brain coupling a useful framework for determining whether hemodynamic correction alters not only the magnitude of brain-behavioral relationships but also their inferred temporal and spatial organization across GCaMP indicators (**Fig.5**).

**Figure 5.**
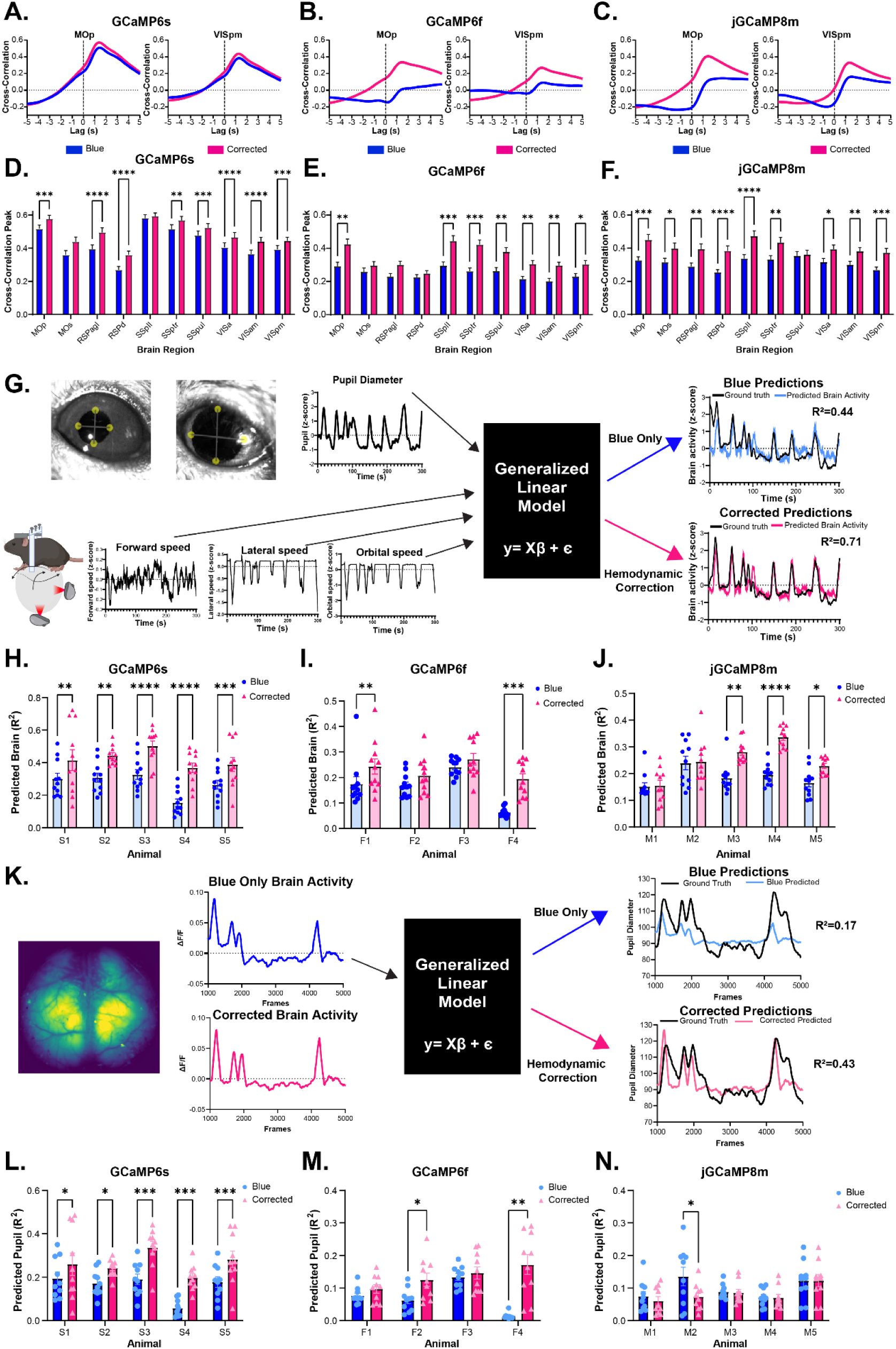
Hemodynamic correction alters pupil-brain coupling and behavioral prediction across GCaMP indicators. **(A-C)** Mean cross-correlation functions between pupil diameter and regional fluorescence signals for representative cortical regions in **(A)** GCaMP6s, **(B)** GCaMP6f, and **(C)** jGCaMP8m mice. For each indicator, MOp is shown on the left and VISpm is on the right. Curves represent averages across 5 animals and 6 recording sessions per animal. Cross correlation is plotted as a function of lag. **(D-F)** Peak cross-correlation values between pupil diameter and regional fluorescence signals across cortical regions for **(D)** GCaMP6s, **(E)** GCaMP6f, and **(F)** jGCaMP8m. Hemodynamic corrected signals showed higher pupil-brain coupling across multiple regions compared with blue only signals. Bars represent mean ± from animals (n=5 per indicator)**. (G)** Schematic of behavior to brain generalized linear modeling. Pupil diameter and airball derived locomotion variables were used as predictors to estimate regional fluorescence activity from either blue only or hemodynamic corrected signals. Model performance was quantified using the coefficient of determination (R^2^) between predicted and observed fluorescence activity. **(H-J)** Behavior to brain model performance across animals for **(H)** GCaMP6s, **(I)** GCaMP6f, and **(J)** jGCaMP8m. Each point represents one cortical region within an animal, and bars indicate mean ± SEM. **(K)** Schematic of reciprocal brain to pupil modeling. Regional blue only or hemodynamic corrected signals were used as predictors to estimate pupil diameter, and model performance was quantified using R^2^ between predicted and observed pupil traces. **(L-N)** Brain to pupil model performance across animals for **(L)** GCaMP6s, **(M)** GCaMP6f, and **(N)** jGCaMP8m. Each point represents one cortical region within an animal, and bars indicate mean ± SEM. Statistical comparisons between blue only and hemodynamic corrected signals were performed using multiple paired two-sided t-test with Holm Šídák correction. Significance is indicated as *p* < 0.05 (*), *p* < 0.01 (**), *p* < 0.001 (***), and *p* < 0.0001 (****).

Cross-correlation analysis revealed strong coupling between pupil dynamics and cortical fluorescence signals across all three GCaMP lines, with region- and indicator-specific temporal profiles. This is shown in representative traces from MOp and VISpm regions **(Fig. 5A-C; Supplementary Fig. 6A-C)**.

These effects were also observed broadly across cortical regions **(Fig. 5D-F)**. Hemodynamic correction significantly increased peak pupil-brain cross-correlation in 8 of 10 regions in GCaMP6s, 7 of 10 regions in GCaMP6f, and 9 of 10 regions in jGCaMP8m following Holm Šídák correction. Although the magnitude of improvement varied across regions and indicators, the overall pattern was remarkably consistent: corrected signals exhibited stronger coupling to pupil dynamics than uncorrected recordings. The broadest enhancement was observed in jGCaMP8m, whereas GCaMP6f exhibited more regionally selective effects **(Fig. 5E-F)**. Together, these results indicate that vascular contributions partially obscure behavior-linked neural activity and that hemodynamic correction strengthens the association between cortical activity and arousal-related behavioral fluctuations across diverse calcium indicators.

We next asked whether stronger coupling was accompanied by changes in temporal organization. Surprisingly, lag at peak cross-correlation remained largely unchanged following correction **(Supplementary Fig. 6D-F)**. Across all GCaMP lines, no cortical region exhibited a significant shift in lag after multiple-comparison correction, and peak correlations remained clustered at positive delays of approximately 1-1.5s. Thus, hemodynamic correction increased the strength of pupil-brain coupling while preserving its temporal organization, indicating that correction enhances behavior-linked signal structure without disrupting the underlying timing relationships between cortical activity and arousal dynamics.

To determine whether these effects extend beyond pairwise correlations, we next evaluated how well behavioral variables predicted cortical activity using generalized linear models (GLM). Pupil diameter and airball-derived locomotion variables were used to predict regional fluorescence signals **(Fig. 5G)**. Across indicators, hemodynamic correction generally improved brain model performance, although the magnitude of improvement differed substantially between GCaMP lines **(Fig. 5H-J)**. The largest gains were observed in GCaMP6s, where corrected signals were substantially more predictable from behavioral variables than blue-only recordings. GCaMP6f exhibited more modest and variable improvements, whereas jGCaMP8m showed smaller overall increases despite retaining significant behavior-related information. These results indicate that removal of vascular contributions can enhance the proportion of cortical variance explained by behavioral state, but that the magnitude of this effect depends on indicator properties.

We next performed the reciprocal analysis by predicting pupil diameter from cortical fluorescence activity **(Fig. 5K)**. This brain-to-pupil model asked whether corrected and uncorrected cortical signals differed in the amount of behaviorally relevant information they contained. Consistent with the behavior-to-brain analysis, GCaMP6s exhibited robust improvements following correction, with significant increases in prediction performance across all animals **(Fig. 5L)**. GCaMP6f showed smaller but generally positive effects **(Fig. 5M)**. In contrast, jGCaMP8m displayed a modest reduction in prediction performance following correction **(Fig. 5N)**. However, baseline prediction performance was substantially lower in jGCaMP8m than in the other indicators, suggesting that correction effects should be interpreted within the context of reduced overall predictability.

Prediction accuracy alone does not reveal which cortical signals drive pupil dynamics. We therefore examined regional model (β) coefficients to determine whether hemodynamic correction altered the cortical substrates contributing to pupil prediction **(Suppl. Fig 6G-I)**. Across all GCaMP lines, corrected and blue-only signals produced distinct coefficients profiles, indicating that correction reorganized the cortical features contributing to pupil prediction rather than simply scaling model performance.

Together, these findings demonstrate that hemodynamic correction reshaped inferred brain-behavior coupling at multiple levels. Correction strengthened pupil-brain cross-correlations without systemically changing temporal lag structure, increased the predictability of cortical activity from behavioral variables, and reorganized the cortical features contributing to pupil dynamics. Importantly, these effects differed across GCaMP indicators, with the strongest and most consistent improvements observed in GCaMP6s. Thus, correction strengthened behavior-linked signal structure while preserving the temporal organization of pupil dynamics.

### Hemodynamic contamination masks glioblastoma-associated changes in cortical dynamics and biases network interpretation

GBM provides a stringent disease-state test of hemodynamic correction because tumor progression directly remodels vascular physiology that contributions to optical signal contamination **(Fig. 6A-B)**. We therefore analyzed blue-only and hemodynamic-corrected signals in parallel across baseline and tumor conditions to determine whether vascular contamination altered the inferred cortical dynamics in disease progression.

**Figure 6.**
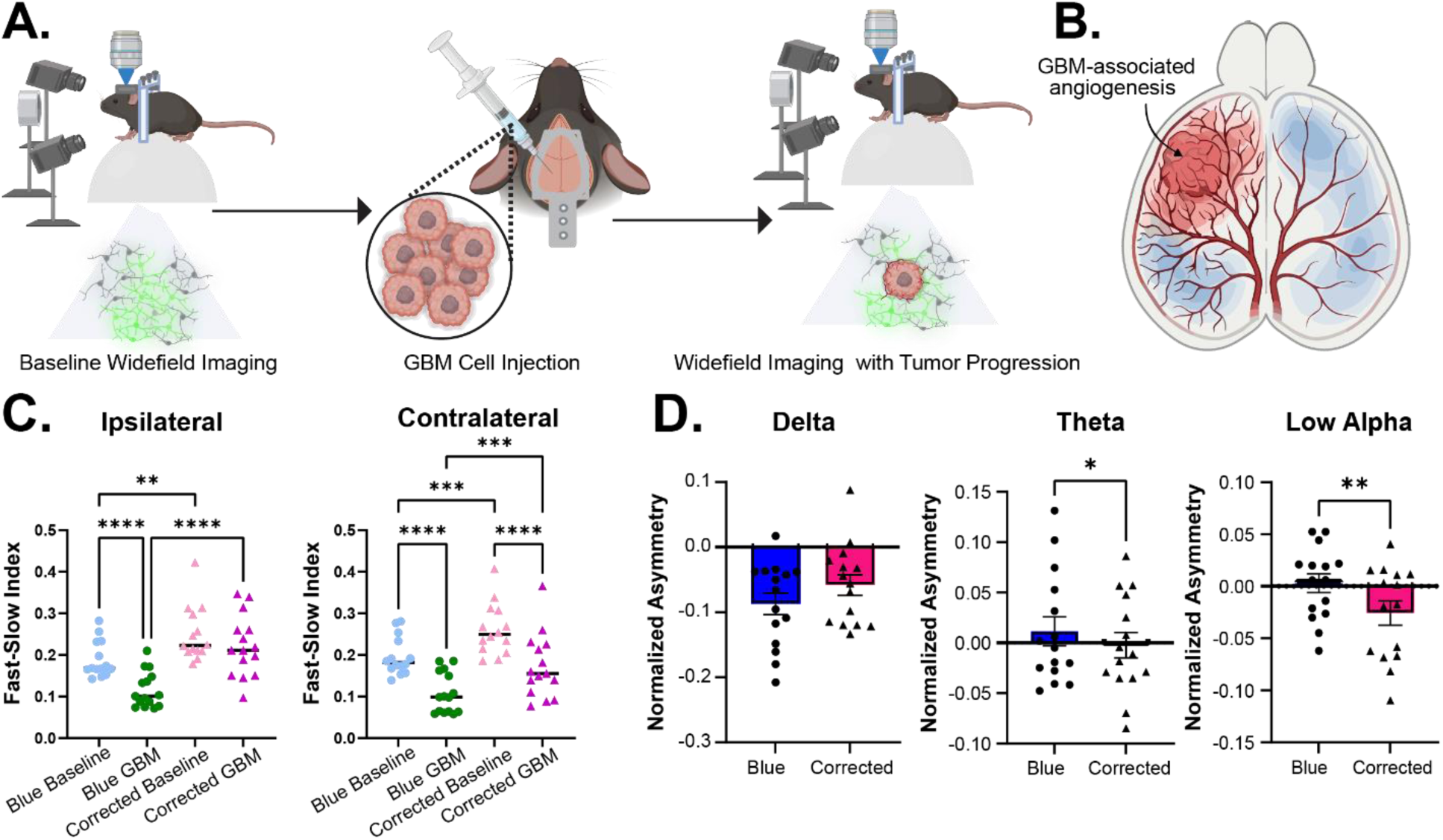
Hemodynamic contributions influence interpretation of cortical dynamics in glioblastoma. **(A)** Experimental schematic showing baseline widefield calcium imaging, intracranial injection of GBM cells and longitudinal imaging following tumor progression. **(B)** Conceptual illustration of GBM associated angiogenesis, highlighting increased vascularization within the tumor bearing hemisphere and its potential contribution to hemodynamic signals that can confound interpretation of neural activity. **(C)** Fast-slow index across baseline and GBM conditions for blue-only and hemodynamic-corrected signals in the ipsilateral and contralateral hemispheres. Hemodynamic correction significantly altered the fast-slow index, with multiple pairwise differences observed between conditions (two-way ANOVA with Tukey’s multiple comparisons test). **(D)** Normalized nodal asymmetry between tumor and contralateral hemispheres across frequency bands (left) delta (*p* = 0.0815), (middle) theta (*p* = 0.0145), and (right) alpha (*p* = 0.0015) bands (paired two-tailed t-tests). Points represent individual animals; bars indicate mean ± SEM. Significance is indicated as *p* < 0.05 (*), *p* < 0.01 (**), *p* < 0.001 (***), and *p* < 0.0001 (****).

In blue-only recordings, GBM shifted cortical activity toward slower dynamics, reflected by reductions in the fast-slow index across both hemispheres **(Fig. 6C)**. Hemodynamic correction changed this interpretation by revealing that the apparent slowing was not uniformly preserved after vascular contributions were removed. Instead, correction exposed a more hemisphere dependent spectral phenotype, indicating that tumor associated vascular fluctuations can make disease effects appear more spatially symmetric in uncorrected signals. Thus, hemodynamic contamination did not simply alter the magnitude of the fast-slow index; it changed the inferred pattern of GBM-associated cortical dysfunction.

To characterize the spatial organization of these effects, node strength was quantified as a measure of regional connectivity with the broader cortical network **(Supplementary Fig. 7A-B)**. Hemodynamic correction revealed region- and hemisphere-specific alterations that were not uniformly apparent in blue-only signals. These spatial differences were further quantified using a normalized asymmetry metric comparing homologous regions between tumor-bearing and contralateral hemispheres **(Fig. 6D; Supplementary Fig. 7C)**. Whereas delta-band asymmetry did not differ significantly between blue-only and corrected signals, theta- and alpha-band asymmetries emerged only after correction, indicating that higher-frequency hemispheric differences are partially masked by vascular contamination in uncorrected recordings.

To determine whether these effects extended beyond static connectivity structure, we next examined time-resolved network dynamics using sliding-window functional connectivity projected into a shared low-dimensional principal component (PC) space **(Fig. 7A)**. Blue-only and corrected signals followed distinct trajectories through network state space, demonstrating that hemodynamic correction reorganizes the temporal evolution of cortical network states rather than simply altering static connectivity strength. Across animals, corrected signals followed more constrained trajectories **(Fig. 7B; Supplementary Fig. 8A-B)**, consistent with broad vascular fluctuations artificially broadening apparent state-space exploration in uncorrected recordings. Mean windowed FC matrices, showed corresponding differences in connectivity structure across regional pairs **(Fig. 7C)**. PC loadings further identified the region-pair features driving the trajectory space, with PC1 and PC2 weighted by distributed motor, retrosplenial, and visual connections **(Fig. 7D)**.

**Figure 7.**
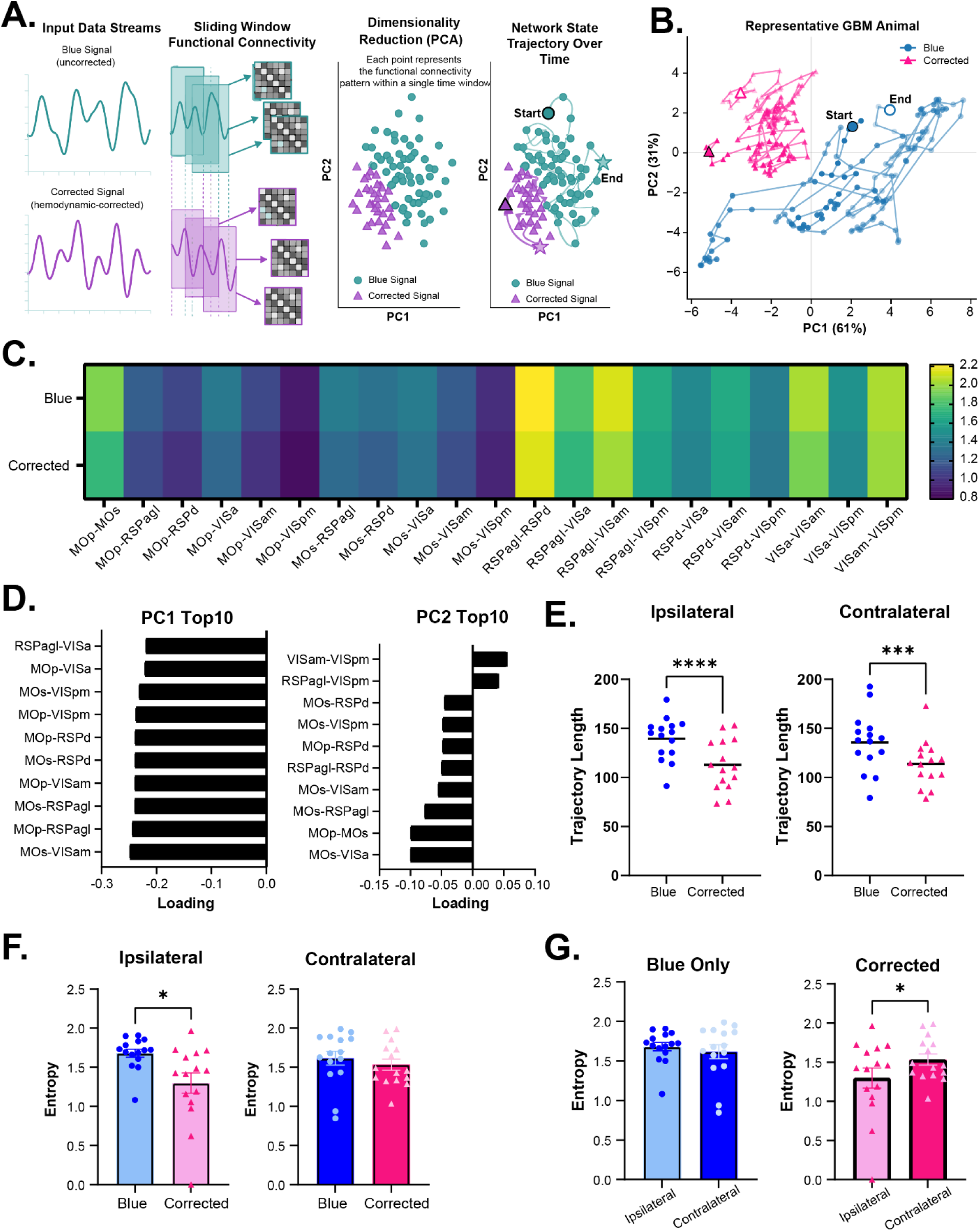
Hemodynamic correction alters dynamic functional connectivity trajectories in GBM. **(A)** Schematic illustrating the analysis pipeline. Fluorescence time series were segmented into sliding windows; functional connectivity matrices were computed for each window and window level FC vectors were projected into PCA space to generate dynamic network trajectories. **(B)** Representative example showing PCA trajectories for a single animal, with blue only signals (blue) and hemodynamic corrected signals (pink), illustrating differences in state space exploration. **(C)** Mean FC matrices over all windows. **(D)** Top contributing region pairs to the first (PC1) and second (PC2) principal components, highlighting features driving differences between signal types. **(E)** Trajectory length in PCA space for (left) ipsilateral and (right) contralateral hemispheres (paired two-tailed t-test; ipsilateral *p* <0.0001, contralateral *p* = 0.0010). **(F)** State entropy for (left) ipsilateral and (right) contralateral hemispheres. Hemodynamic correction significantly altered entropy in ipsilateral (*p* = 0.0107) but not contralateral (*p* = 0.2487), indicating region specific changes in network state diversity. **(G)** Comparison of hemispheric differences in entropy for (left) blue only (p =0.4824) and (right) corrected signals (*p* = 0.0343), indicating correction reveals hemispheric asymmetries that are obscured in uncorrected data. Points represent individual animals; bars indicate mean ± SEM. Significance is indicated as *p* < 0.05 (*), *p* < 0.01 (**), *p* < 0.001 (***), and *p* < 0.0001 (****).

These differences were reflected in the geometry of the trajectories themselves. Blue-only signals occupied broader regions of state space, whereas corrected signals followed more spatially constrained paths. Consistent with this observation, trajectory length was significantly reduced following correction in both the ipsilateral (*p* < 0.0001) and contralateral hemispheres (*p* = 0.0010) **(Fig. 7E)**, indicating that uncorrected signals overestimate the extent of network-state exploration.

Changes in trajectory organization were accompanied by altered in network-state diversity. State entropy, calculated as the Shannon entropy of state occupancy over time, was used to quantify the diversity and flexibility of network dynamics, with higher values indicating greater variability in state transitions. Hemodynamic correction significantly reduced entropy within the ipsilateral hemisphere, but not in the contralateral hemisphere **(Fig. 7F)**, showing that hemodynamic contamination disproportionally inflates apparent network variability within tumor-affected cortex and obscures disease-associated constraints on cortical dynamics. The number of distinct states visited and the frequency of transitions between states were also significantly altered in the ipsilateral hemisphere, but not in the contralateral hemisphere **(Supplementary Fig. 8C-D)**.

Importantly, these effects fundamentally altered the biological interpretation of hemispheric differences in GBM. In blue-only signal, we observed no significant differences between ipsilateral and contralateral hemispheres, demonstrating comparable levels of network variability across hemispheres regardless of tumor burden. In contrast, hemodynamic correction revealed significantly reduced entropy within the tumor bearing hemisphere **(Fig. 7G),** indicating that GBM-associated cortical dynamics are more constrained and less diverse than would be inferred from uncorrected signals. Thus, hemodynamic contamination can mask disease-related alterations in network-state organization and alter conclusions regarding the functional impact of tumor progression.

To determine whether corrected and uncorrected signals occupied distinct regions of functional connectivity space, we performed multivariate classification analysis using linear discriminant analysis (LDA). This analysis demonstrated clear separation between blue-only and corrected recordings within functional connectivity space (**Supplementary Fig. 8E-F),** identifying consistent connectivity patterns that reliably distinguished the two conditions with classification accuracy well above chance (mean ± SEM: 0.965 ± 0.018 for blue-only and 0.954 ± 0.023 for corrected). These results indicate that the observed differences reflect systematic organization of network-state representations in GBM rather than variability within the dataset.

Across spectral, hemispheric, and time-resolved connectivity analyses, the hemodynamic correction changed the inferred GBM-associated cortical phenotype. Blue-only signals emphasized slower dynamics and larger apparent state-space exploration, whereas corrected signals revealed frequency specific hemispheric asymmetry and more constrained network state dynamics, particularly in the tumor bearing hemisphere. Thus, vascular contamination altered both the magnitude and the spatial pattern of GBM-associated network changes detected from widefield recordings.

### Deep learning prediction of hemodynamically corrected signals recovers spectral and network organization across laboratories and disease states

The preceding analyses showed that hemodynamic correction can substantially alter the inferred organization of cortical activity across spectra, network, behavioral, and disease context. However, many existing widefield datasets contain only blue-channel recordings, limiting the ability to apply dual-wavelength correction retrospectively. To address this challenge, we developed HemoTCN (**Supplementary Video 1**), an open-source deep learning framework that predicts hemodynamically corrected fluorescence directly from blue-channel widefield recordings (**Fig. 8A**).

**Figure 8.**
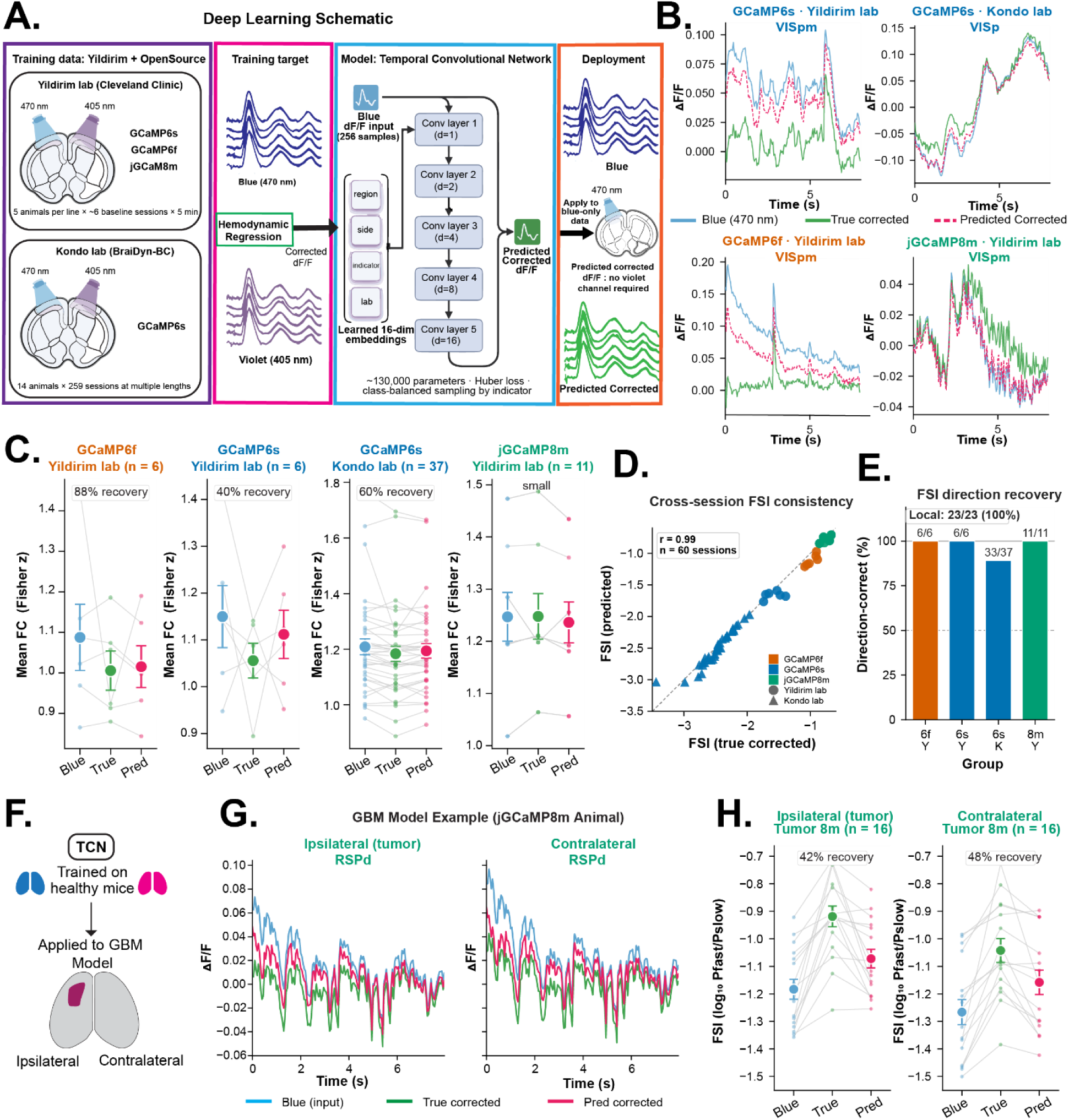
Deep learning prediction of hemodynamic corrected signals using HemoTCN. **(A)** Schematic of the HemoTCN deep learning workflow. Dual wavelength recordings from multiple labs containing blue only and ground truth corrected fluorescence signals were used to train a residual temporal convolutional network to predict hemodynamic corrected traces directly from blue channel input without simultaneous violet channel acquisition. **(B)** Representative fluorescence traces from multiple animals showing blue, ground truth corrected (green), and model predicted signals (pink dashed). **(C)** Functional connectivity (FC) recovery across labs. Recovery percentage quantifies the extent to which the model recapitulates the shift from blue only to true corrected connectivity. GCaMP6f recordings exhibited the strongest recovery (∼88%), while GCaMP6s from Yildirim and Kondo datasets shows ∼40% and ∼60%, respectively. jGCaMP8m exhibited small FC shifts overall, consistent with reduced hemodynamic contamination. **(D)** Fast slow index (FSI) consistency between true corrected and predicted corrected signals across sessions (Pearson correlation r = 0.99). **(E)** FSI direction recovery showing whether the model correctly predicts the direction of FSI change relative to blue only signals. Direction recovery reached 100% for Yildirim lab recordings and 33/37 sessions for Kondo dataset. **(F)** Schematic illustrating out of distribution application of HemoTCN in a disease model, glioblastoma (GBM) recordings without retraining or fine-tuning. **(G)** Representative ipsilateral and contralateral GBM fluorescence traces showing blue only (blue), true corrected (green), and predicted corrected (pink) signals. **(H)** FSI recovery in GBM recordings for ipsilateral and contralateral hemispheres, demonstrating that predicted signals closely approximate true corrected signals despite the model never being trained on tumor data.

HemoTCN is based on residual temporal convolutional networks trained to learn correction-relevant structure from blue-channel activity alone. The model was trained using 357,500 temporal windows spanning 23 cortical regions, three GCaMP indicators, multiple imaging configurations, and two independent laboratories. This design was intended to promote generalization across experimental conditions rather than optimization for a single dataset. In addition to inference, the HemoTCN platform includes graphical user interface (GUI)-based tools for correction quality assessment, transfer learning, laboratory-specific fine-tuning, and de-novo model training **(Supplementary Fig. 9).**

Across held out recordings, HemoTCN accurately reconstructed the temporal structure of dual-wavelength corrected signals **(Fig. 8B)**. Predicted signals closely matched experimentally corrected fluorescence traces, achieving a held-out Pearson correlation of 0.971 and mean absolute error of 0.00498. These results indicate that the model captures correction-relevant temporal structure rather than simply smoothing or denoising fluorescence recordings.

We next asked whether HemoTCN could recover the higher-order network features shown throughout this study to be sensitive to hemodynamic contamination. Functional connectivity analyses demonstrated that predicted signals reproduced correction-dependent reductions in large-scale cortical synchrony across multiple GCaMP indicators and laboratories **(Fig. 8C)**. Recovery was strongest in GCaMP6f, where predicted signals closely approximated the connectivity shift observed following dual-wavelength correction. Similar trends were observed in GCaMP6s signals acquired in both the Yildirim and Kondo et al. datasets, demonstrating generalization across independent experimental environments. Importantly, in jGCaMP8m recordings, where blue-only and corrected signals were already highly similar, HemoTCN did not introduce large artificial changes, indicating that the model preserves underlying signal structure when correction effects are modest.

Because spectral analyses represented a major source of biological reinterpretation throughout the manuscript, we next evaluated recovery of the fast-slow index (FSI), a summary metric of the balance between higher-frequency neural activity and slow hemodynamic-dominated power. Across healthy test sessions, predicted and experimentally corrected FSI values exhibited near-perfect agreement (r = 0.993; **Fig. 8D**). HemoTCN consistently recovered the direction and magnitude of correction-induced spectral shifts across GCaMP indicators and laboratories **(Fig. 8E),** demonstrating robust reconstruction of spectral features that are strongly influenced by vascular contamination. Together, these findings indicate that HemoTCN recovers correction-dependent spectral organization across both local and publicly available datasets.

To further evaluate model robustness, we applied HemoTCN to GBM recordings without retraining or fine-tuning **(Fig. 8F)**. This represents a stringent out-of-distribution test because the model was trained exclusively on healthy recordings and therefore had not previously encountered tumor-associated vascular remodeling. Despite this challenge, HemoTCN consistently shifted spectral metrics toward the experimentally corrected state in both ipsilateral and contralateral hemispheres **(Fig. 8G-H)**. Although prediction performance was reduced relative to healthy recordings, the model recovered a substantial fraction of the correction-induced spectral shift, demonstrating meaningful generalization to disease-associated vascular abnormalities.

Beyond predictive performance, HemoTCN was designed as a practical framework for extending hemodynamic correction to laboratories lacking custom dual-wavelength systems **(Supplementary Fig. 9)**. The software platform supports blue-only inference, correction quality evaluation, transfer learning, model fine-tuning, and training of laboratory-specific models. Integrated visualization tools allow users to compare corrected and uncorrected signals using power spectral density, fast-slow index (FSI), band power distributions, and functional connectivity analyses. These features enable researchers to apply pretrained models to legacy datasets, adapt correction models to their own recordings, or develop customized models when paired dual-wavelength data are available.

Together, these results demonstrate that HemoTCN learns biologically meaningful correction-relevant structure from blue-channel recordings and recovers key network and spectral signatures of dual-wavelength hemodynamic correction. Importantly, the model generalizes across laboratories, calcium indicators, and disease states, suggesting that deep learning can provide a scalable strategy for extending hemodynamic correction to the large collection of existing blue-only widefield imaging datasets. By enabling retrospective correction of previously inaccessible recordings, HemoTCN broadens the practical impact of hemodynamic correction and facilitates more accurate inference of cortical dynamics across experimental settings.

## Discussion

Widefield calcium imaging has become a central tool for studying cortex-wide neural dynamics, yet interpretation of fluorescence signals remains fundamentally complicated by vascular contributions embedded within optical recordings. Here, we demonstrate that hemodynamic correction does not simply rescale fluorescence measurements but systematically reshapes inference of cortical organization across spatial, spectral, behavioral, and disease-related domains. Across GCaMP indicators, correction altered functional parcellation, reduced large-scale connectivity, increased network segregation, redistributed spectral power, strengthened brain-behavior coupling, and changed the interpretation of tumor-associated cortical dynamics. Together, these findings extend previous work demonstrating substantial hemodynamic contributions to widefield fluorescence measurements^2,20,22,39–41^ and establish that vascular signals fundamentally influence how mesoscale brain activity is interpreted.

A central conclusion emerging from this work is that hemodynamic contamination behaves as a structured biological confound rather than random measurement noise. The problem is not simply that vascular variance is present, but that it is organized along similar covariances, frequencies, and state related dimensions used to infer mesoscale cortical dynamics in widefield calcium imaging. Measures such as functional connectivity, spectral power, network modularity, dimensionality reduction, and brain-behavior coupling all depend on shared temporal structure. Therefore, vascular components change not merely the amplitude of individual measurements, but the relationships among cortical regions and associated features. In our results, this shifted the outcomes across multiple analytical domains and, in some cases, altered the biological conclusions drawn from the same dataset (**Fig. 9**).

**Figure 9.**
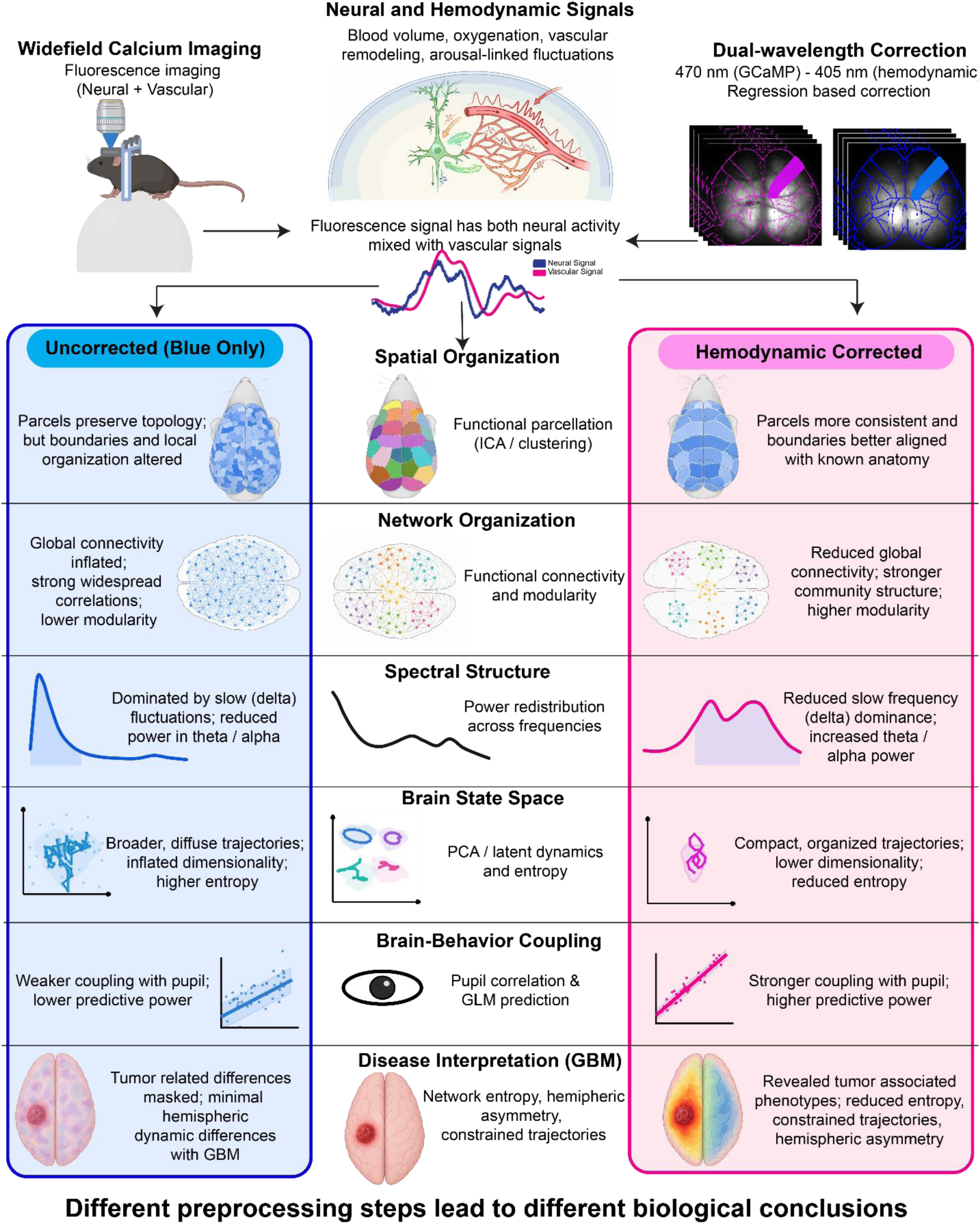
Conceptual framework illustrating how structured hemodynamic signals reshape inference of cortical dynamics in widefield calcium imaging. Widefield calcium imaging records fluorescence signals that contain both neuronal and vascular contributions. Structured hemodynamic signals arising from blood volume fluctuations, oxygenation changes, vascular remodeling, and arousal-related physiology propagate through multiple stages of analysis and influence interpretation of mesoscale brain activity. In uncorrected (blue-only) recordings, hemodynamic contributions can alter functional parcellation, inflate functional connectivity, reduce network modularity, bias spectral structure toward slow-frequency activity, distort low-dimensional brain-state representations, weaken inferred brain–behavior relationships, and obscure disease-associated phenotypes. Dual-wavelength hemodynamic correction separates vascular and neuronal contributions, revealing more refined spatial organization, reduced global synchrony, increased network segregation, enhanced representation of faster neural dynamics, stronger brain–behavior coupling, and previously masked disease-related signatures. In glioblastoma (GBM), correction exposes hemispheric asymmetries, reduced network entropy, and constrained state-space trajectories that are attenuated or obscured in uncorrected recordings. Together, these findings demonstrate that hemodynamic contamination is not merely a source of measurement noise but a structured biological signal that systematically influences inference of cortical organization, network dynamics, and disease-associated phenotypes. HemoTCN extends these principles to blue-only datasets by enabling scalable prediction of hemodynamically corrected signals from fluorescence recordings alone.

Importantly, our goal is not to claim that hemodynamic-corrected signals represent an absolute ground truth for neuronal population activity. Rather, our findings demonstrate that biological conclusions depend strongly on how vascular contributions are handled. The key implication is therefore not that one signal is definitively correct, but that vascular structure must be explicitly considered when interpreting mesoscale fluorescence recordings.

### Hemodynamic contamination reshapes inference of cortical organization and network structure

One of the earliest consequences of hemodynamic correction emerged at the level of functional organization. Independent component analysis revealed that corrected and uncorrected recordings often preserved broad cortical topology while substantially reorganizing local parcel boundaries. This distinction is important because many widefield studies rely on data-driven parcellation approaches to define functional units prior to downstream analyses^4,32,42^. In these settings, hemodynamic contamination does not simply influence the activity measured within a predefined region; it can influence how regions themselves are identified. Thus, correction can shift the functional units that enter downstream analyses, altering the spatial foundation from which network structure and brain-behavior relationships are inferred.

The magnitude of these effects depended on both cortical location and calcium indicator. We showed that GCaMP6f consistently exhibited the largest variance removed fractions across atlas-based and pixel-based analyses, indicating a stronger overlap between fluorescence fluctuations and the hemodynamic reference signal. Importantly, this should not be interpreted as evidence that one indicator is inherently superior or inferior. The magnitude of correction likely reflects a complex interaction among indicator kinetics, baseline brightness, expression patterns, signal-to-noise properties, and the relative contribution of neural and vascular variance ^43–46^. Our findings therefore suggest that hemodynamic correction cannot be assumed to have uniform consequences across indicators. Instead, indicator selection and correction strategy should be considered jointly during experimental design, as sensor properties influence not only which neural dynamics can be measured but also the extent to which downstream analyses are affected by vascular contamination^43,47^.

The effects of hemodynamic contamination became particularly evident at the network level. Our functional connectivity analyses revealed that blue-only recordings consistently exhibited stronger cortex-wide synchrony than corrected signals, whereas corrected networks displayed increased modularity and more clearly defined community structure. These observations suggest that spatially widespread vascular fluctuations introduce shared variance across cortical regions, artificially inflating estimates of inter-regional coupling. Because correlation-based connectivity measures are highly sensitive to common fluctuations, even modest vascular contributions can increase apparent synchrony without reflecting underlying neural interactions. Removal of this shared variance reduced global connectivity and increased network segregation, providing a mechanistic explanation for the observed increase in modularity. These findings have important implications for studies using connectivity-based approaches to investigate cortical organization during behavior, development, injury, and disease^1,4,5,7,20–22,40,41^.

Spectral analyses provided a complementary perspective on these network-level effects. Across indicators, we demonstrated that hemodynamic correction consistently reduced slow-frequency dominance and increased the relative contribution of theta- and alpha-band activity, indicating that uncorrected fluorescence signals are biased toward slower dynamics. This observation is consistent with prior studies showing that hemodynamic contamination contributes substantially to low-frequency fluctuations in widefield recordings^20,22^. Many physiological processes including blood volume changes, oxygenation dynamics, vasomotion, respiration-linked oscillations, and systemic cardiovascular fluctuations occur within frequency ranges commonly analyzed in optical imaging^38,40^. Consequently, slow spectral power in uncorrected recordings may reflect a mixture of vascular and neural processes rather than neuronal activity alone. The increase in fast-slow index following correction supports this interpretation and suggests that slow vascular variance can mask or compress faster calcium-associated dynamics.

Importantly, the effects of correction were not limited to individual analytical metrics. Our multivariate analyses demonstrated that hemodynamic correction produced coordinated shifts across connectivity, spectral structure, and network topology, resulting in distinct low-dimensional representations of cortical state space. Rather than acting independently on isolated measurements, correction systematically reorganized the latent structure of the dataset itself. This observation is particularly significant because biological interpretations are rarely based on a single metric. Modern systems neuroscience studies often integrate connectivity, oscillatory structure, network organization, and behavioral variables to characterize cortical states. Our findings therefore suggest that hemodynamic correction changes the analytical state of a dataset rather than simply modifying the amplitude of individual fluorescence traces.

### Hemodynamic correction reveals stronger and more interpretable brain–behavior relationships

The brain-behavior coupling analyses provided an important counterpoint to the connectivity findings and highlighted the complexity of hemodynamic influences on cortical recordings. Based on the strong association between pupil diameter, arousal, autonomic physiology, and vascular state, we initially hypothesized that blue-only signals would exhibit stronger coupling to pupil dynamics. Surprisingly, the opposite pattern emerged. Across indicators, hemodynamic correction frequently strengthened pupil-brain correlations and improved predictive modeling performance, indicating that vascular contamination can obscure behaviorally relevant neural signals rather than simply inflate them.

These findings support a more nuanced interpretation of the relationship between arousal, neural activity, and hemodynamics. Although pupil diameter is closely linked to vascular physiology, it is also a robust marker of low-dimensional brain state and neuromodulatory activity^19,48^. Widefield one-photon imaging does not resolve individual neurons but instead measures population-level fluorescence arising from large neural ensembles distributed across the dorsal cortex^3,49^. At this mesoscale level, pupil fluctuations likely reflect coordinated cortical state transitions rather than isolated local events, consistent with previous studies demonstrating that pupil dynamics track large-scale neural activity, cortical state changes, and neuromodulatory signaling^27–29^. Pupil dynamics can also provide a low-dimensional representation of spatiotemporal brain activity that incorporates neural, metabolic, and blood-oxygen signals ^50^. Within this framework, hemodynamic correction may strengthen pupil-brain coupling because it reduces vascular distortions that interfere with neural population structure associated with arousal and state regulation.

Our GLM analyses further extend this conclusion by demonstrating that correction influences both prediction strength and inferred cortical substrates of behavior. Behavior-to-brain models generally performed better following correction, particularly in GCaMP6s, indicating that pupil and locomotion explained a greater proportion of variance in corrected signals. In reciprocal brain-to-pupil models, correction improved prediction performance in GCaMP6s and GCaMP6f, but not in jGCaMP8m, where overall predictive power was lower. This reduced predictability may reflect diminished slow-state content, faster indicator kinetics, or limitations imposed by the 20 Hz acquisition rate. Importantly, our coefficient analyses demonstrated that correction altered which cortical regions contributed to behavioral prediction, indicating that hemodynamic correction changes not only the strength of brain–behavior relationships but also the inferred cortical architecture supporting those relationships.

### Disease-associated vascular remodeling amplifies interpretational bias in glioblastoma

The implications of these findings become particularly important in disease states where vascular dysfunction is itself a defining feature of pathology. GBMs are characterized by extensive angiogenesis, abnormal vascular remodeling, blood-brain barrier disruption, and altered neurovascular coupling, creating a tumor microenvironment in which neural and vascular signals become increasingly difficult to disentangle^34–36^. Under these conditions, hemodynamic contamination is not simply superimposed onto neural activity but becomes intertwined with the biological processes under investigation. Consequently, accurate interpretation of tumor-associated cortical dynamics requires consideration of both neural and vascular contributions to the measured fluorescence signal.

Our GBM analyses demonstrate that hemodynamic correction can fundamentally alter biological conclusions. In the spectral domain, blue-only recordings suggested a pronounced shift toward slower cortical dynamics during tumor progression. Following correction, however, the relationship between baseline and tumor states was substantially reorganized, indicating that a portion of the apparent slowing reflected tumor-associated vascular contributions rather than neural activity alone. Similarly, correction revealed frequency-specific hemispheric asymmetries that were not evident in uncorrected recordings, demonstrating that interpretation of tumor-associated lateralization depends strongly on whether vascular contributions are retained or removed.

The most striking effects emerged in the analysis of dynamic network organization. Uncorrected signals occupied broader regions of network state space, exhibited longer trajectories, and displayed higher apparent entropy, suggesting extensive network variability and flexibility. Following correction, network trajectories became more constrained and organized, particularly within the tumor-bearing hemisphere. Importantly, blue-only recordings suggested little difference in network-state diversity between hemispheres, whereas corrected signals revealed significantly reduced entropy ipsilateral to the tumor. This distinction fundamentally changes the biological interpretation of glioblastoma-associated network dysfunction. Rather than concluding that tumor progression produces minimal hemispheric differences in cortical dynamics, correction revealed a previously obscured phenotype characterized by constrained state-space exploration and reduced network flexibility within tumor-affected cortex. These findings illustrate how vascular contamination can mask disease-associated neural signatures and, in some cases, determine whether a phenotype is detected at all.

More broadly, these observations parallel challenges encountered in human functional neuroimaging. Previous studies have shown that glioma progression can produce both local and distributed connectivity alterations, with vascular dysfunction contributing to changes in functional organization^51^. Clinical fMRI faces a related challenge in which tumor-induced neurovascular uncoupling can attenuate or eliminate BOLD responses despite preserved neural function, producing false-negative activation maps and misleading interpretations of cortical reorganization during tumor progression^52,53^. Our findings suggest that a similar problem exists in widefield calcium imaging. Tumor-associated vascular remodeling can generate structured fluctuations that are easily misinterpreted as neural dynamics, while genuine neural alterations may be obscured by vascular variance.

### Deep learning enables scalable correction of legacy widefield imaging datasets

Because our results demonstrate that hemodynamic correction can substantially reshape biological interpretation, scalable correction approaches are needed for the large number of existing datasets acquired without dual-wavelength imaging. To address this challenge, we developed HemoTCN, a deep learning framework that predicts hemodynamically corrected signals directly from blue-channel recordings.

Importantly, HemoTCN was designed not merely to reproduce fluorescence traces, but to recover the network-level and spectral features shown throughout this study to be sensitive to vascular contamination. The model generalized across laboratories, calcium indicators, and imaging configurations, and retained meaningful predictive performance when applied to glioblastoma recordings that were never encountered during training. These findings support the possibility of extending correction to historical datasets that would otherwise remain inaccessible to modern correction approaches.

### Limitations and future directions

Several limitations should be considered. First, the dual-wavelength correction approach used here relies on a 405 nm reference channel, which provides an effective but imperfect estimate of hemodynamic contributions. This approach does not directly measure oxyhemoglobin and deoxyhemoglobin dynamics, nor does it fully account for nonlinear absorption, scattering, or pathlength effects. Future multispectral approaches may provide more physiologically specific correction^20–22,54^.

Second, the 20 Hz effective sampling rate limits characterization of faster neural dynamics, particularly for indicators such as jGCaMP8m whose kinetics are optimized for rapid calcium transients. Third, although dual-wavelength correction provides a principled strategy for reducing vascular contamination, the present study does not establish an absolute ground truth for neuronal population activity. Simultaneous electrophysiology, voltage imaging, or additional multimodal approaches will be necessary to determine which corrected features most accurately reflect underlying neural dynamics and state transitions.

Future studies integrating electrophysiology, multispectral imaging, and computational modeling may further clarify how vascular and neural signals interact to shape large-scale cortical dynamics. Extending similar analyses to additional disease models, including neurodegeneration, stroke, and neurodevelopmental disorders, may reveal whether vascular contributions similarly influence systems-level interpretations across diverse neurological conditions.

### Conclusions

In conclusion, our findings demonstrate that hemodynamic correction is not merely a preprocessing choice but a critical determinant of biological interpretation in mesoscale calcium imaging. Across healthy and glioblastoma-bearing mice, vascular contributions systematically reshaped inferred cortical organization, network architecture, spectral structure, brain-behavior relationships, and low-dimensional state-space dynamics. These effects propagated across analytical domains and, in several cases, altered whether disease-associated phenotypes were detected at all. By integrating dual-wavelength imaging, multiscale network analysis, disease-state evaluation, and deep learning-based correction, this work establishes hemodynamic correction as a central component of experimental design, analysis, and interpretation in widefield calcium imaging. More broadly, our findings suggest that vascular physiology should be considered an integral component of system-level optical imaging experiments, as decisions regarding hemodynamic correction can influence not only measured activity but the biological conclusions derived from it.

## Methods

### Animals and Experimental Design

All experimental procedures were approved by and performed in accordance with guidelines established by the Cleveland Clinic Research Institute Institutional Animal Care and Use Committee (IACUC). All parental lines were purchased from Jackson Laboratories and subsequently bred in house to generate experimental animals. Mice were ad libitum access to food and water.

Experimental imaging animals were bred by crossing a CaMKIIa driver line with a Cre-dependent GCaMP reporter line to generate mice with expression of genetically encoded calcium indicators in cortical excitatory neurons. B6.Cg-Tg(CaMK2a-cre)T29-1Stl (Jackson Laboratory, stock #005359) was bred with one of the following reporter lines: Ai96(RCL-GCaMP6s) (Jackson Laboratory, stock #024106), Ai148D (TIT2-GC6f-ICL-tTA2)-D (Jackson Laboratory, stock #030328), or TIGRE2-jGCaMP8m-IRES-tTA2-WPRE (Jackson Laboratory, stock #037718). Offspring genotypes were confirmed using PCR based genotyping services (Transnetyx, Memphis, TN).

Widefield imaging data were collected from three GCaMP expressing mouse lines (GCaMP6s, GCaMP6f, and jGCaMP8m), with five animals per group (n=5 per group, 15 total). Each animal was recorded across six independent sessions, resulting in repeated measurements used for within animal averaging and group-level comparisons.

GBM experiments were performed in jGCaMP8m mice (n=16) using stereotaxic implantation of syngeneic SB28 tumor cell line. SB28 cells carrying NRasV12/Pt2/Shp53/mPDGF alterations were used. A total of 20,000 cells were pressure injected unilaterally into the left hemisphere using a 31-gauge needle mounted onto a stereotaxic frame (RWD #68535) at A.P +1.2 mm, M.L. −1.6 mm, D.V. −3.8 mm relative to bregma. Following injection, the craniotomy was sealed with bone wax and animals recovered under standard postoperative care. Animals were imaged at baseline and again near endpoint following tumor progression. For hemisphere specific analyses, the left hemisphere was defined as ipsilateral/tumor-bearing and the right as contralateral.

### Surgical Procedures

All surgical procedures were performed under isoflurane anesthesia (2.5% induction; 1-1.5% maintenance), with animal’s temperature maintained on a feedback-controlled heating pad (ThermoStar Homeothermic Monitoring System, RWD). The scalp was shaved (Vetiva Mini Hair Trimmer, Wahl) and remaining hair was removed (Nair). The scalp was then sterilized using alternating scrubs of betadine and 70% ethanol. Local anesthetic (bupivacaine, 5 mg/kg) was administered, and ophthalmic lubricant (Optixcare) was applied to prevent corneal drying. Pre-operative analgesia and supportive care included buprenorphine XR (3.25 mg/kg), meloxicam (5mg/kg), and sterile saline (0.9% NaCl, 0.5mL/g), delivered subcutaneously.

Animals were secured in a stereotaxic frame, and a midline incision was made to expose the skull. The periosteum and residual membranes were removed to create a clean, dry surface. The skin edges were affixed to the skull using cyanoacrylate adhesive (VetBond, World Precision Instruments), followed by application of a thin base layer of dental cement (C&B Metabond, Parkell). A custom titanium headplate (eMachineShop, NJ) was centered over the dorsal cortex. Additional cement was applied to reinforce the headplate and ensure a stable bond.

A transparent cranial window was forced by applying multiple thin layers of UV-curable optical adhesive (NOA 81, Norland Inc.) over the exposed skull, with each layer cured using 365 nm UV light source (LIGHTFE, UV301 Plus). A 3D printed light isolation cone was secured to the headplate with dental cement to minimize ambient light contamination.

Following surgery, animals were removed from the stereotaxic frame and allowed to recover on a warming pad until fully ambulatory. Post-operative care included meloxicam (5 mg/kg) and sterile saline administration for at least 3 days.

### Widefield One-Photon Dual Wavelength Optical Imaging and Combined Behavioral Recordings

Recordings used systems previously described^55^. Cortex wide fluorescence imaging was performed using a dual-wavelength, sequential illumination workflow in which excitation (470 nm; M470L5, Thorlabs) frames were interleaved with reference (405 nm; M4055LP1, Thorlabs) frames (∼1 ms delay). Excitation paths were combined using a dichroic beamsplitter (87-063, Edmund Optics) and projected onto the dorsal cortical surface through an objective lens (MVL50M23, Navitar; f=50mm, F2/8). Emitted fluorescence was separated using an emission dichroic (T495LPXR, Chroma) and further filtered with a bandpass filter (86-963, Edmund Optics), then collected through a secondary objective (MVL16M23, Navitar, f=16mm, F1.8) and imaged onto a CMOS camera (CS505MU1, Kiralux).

Images were acquired at 1200 x 1200 pixels with 4 x 4 on chip binning (effective resolution: 300 x 300 pixels), corresponding to ∼9.85 x 9.85 mm^2^ cortical field of view. Imaging was performed at 40 Hz with alternating excitation wavelengths, yielding an effective sampling rate of 20 Hz per channel. Delivered optical power was maintained below 10 mW (470 nm) and 50 mW (405 nm).

Behavioral signals were acquired simultaneously with pupil imaging. Pupil dynamics were recorded at 40 Hz using infrared illumination and a laterally positioned camera. Locomotion was measured using spherical treadmill, where a Styrofoam ball was supported on pressurized air. Optical sensors positioned orthogonally around the ball tracked displacement, enabling extraction of forward (translational), lateral (orbital), and rotational (yaw) velocity components, providing frame synchronized behavioral measurements.

### Image Post-processing

Imaged data were acquired as sequential TIFF stacks and concatenated into a single continuous recording for each session. Interleaved wavelength frames were separated into channel specific image stacks based on frame index using a custom ImageJ macro. To remove acquisition transients and standardize the starting point across recordings, the first frame of each separated stack was discarded prior to further processing.

### Allen Atlas Based Hemodynamic Regression

Recordings were rotated to align to the cortical midline and registered to the Allen Mouse Brain Common Coordinate Framework (CCFv3) using landmark based non-reflective similarity transformations. The warped atlas was used to assign pixels to anatomical parcels and extract region-averaged fluorescence time series from both channels.

Following atlas registration, region averaged fluorescence time series were extracted. For each session, stacks were treated as 3D volumes and regional signals were computed by averaging pixel intensities within binary masks for predefined atlas regions. Analysis focused on 10 cortical brain regions spanning motor (MOp, MOs), retrosplenial (RSPagl, RSPd), somatosensory (SSP-ll, SSp-tr, SSP-ul), and visual (VISa, VISam, VISpm).

Hemodynamic contamination was corrected using a region-wise regression approach in which the 405 m signal served as a hemodynamic reference for the 470 nm fluorescence signal. For each region, an ordinary least squares linear model with intercept was fit:

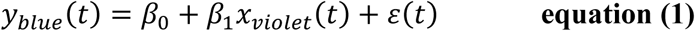

where *y_blue_*(*t*) is the measured calcium-dependent signal and *x_violet_*(*t*) represents hemodynamic fluctuations. The hemodynamic contribution was estimated as the fitted component *ŷ*(*t*), and the corrected signal was defined as the residual:

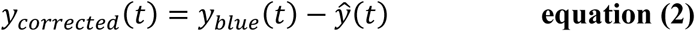

Missing samples were interpolated using a nearest-neighbor approach to ensure consistent signal lengths prior to regression.

Fluorescence signals were normalized using a sliding window median baseline:

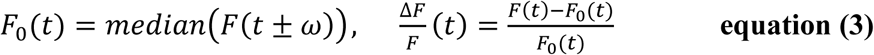

### Pixel Based Hemodynamic Regression

In addition to the atlas-based analysis, hemodynamic correction was performed at the pixel level to capture spatially heterogeneous vascular contributions. For each recording, filtered 470 nm and 405 nm image stacks were treated as time-resolved signals at each pixel location within a defined mask to exclude non-brain pixels. For each pixel, an ordinary least squares linear model similar to that used in the atlas based was used to predict the blue signal from violet reference. Model fitting was performed independently for each pixel using only finite-valued samples. To improve spatial stability, regression coefficients were optionally smoothed using a Gaussian kernel prior to reconstruction of the corrected signal.

### Variance Removed Fraction

The contribution of hemodynamic signals to the observed fluorescence was quantified using the variance removed fraction (VRF) following regression-based correction. VRF was calculated from the regression corrected signals as:

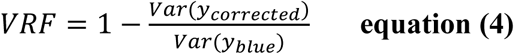

where *y_blue_* is the original fluorescence signal and *y_corrected_* is the hemodynamic regression. This metric represents the proportion of variance in the original signal explained and removed by the hemodynamic component.

For atlas-based analyses, VRF was computed from region averaged fluorescence time series. For pixel-based analyses, VRF was first computed at the pixel level and subsequently averaged within atlas-defined regions to enable direct comparison across methods. In addition, VRF was evaluated as a function of parcel stability by grouping regions according to the matched-component spatial similarity.

All VRF measurements were computed at the session level and averaged across sessions to obtain a single representation value per animal for group level comparisons.

### PCA/ICA Functional Parcellation and Stability Classifications

To generate data-drive functional regions, group level independent component analysis (ICA) was performed on pixel level fluorescence data. A common cortical mask was constructed across sessions by retaining pixels that were classified as brain in at least 80% of recording, ensuring a consistent set of pixels for analysis. For each session, temporally filtered image stacks were reshaped into pixel wise series and standardized by z-scoring each pixel over time.

Dimensionality reduction was first performed using incremental principal component analysis (PCA) followed by ICA to extract spatially independent components. Spatial component maps were reconstructed into image space and converted into discrete parcels using a winner take all assignment where each pixel was assigned to the component with the highest absolute loading.

To compare functional organization before and after hemodynamic correction, spatial correspondence between ICA components derived from uncorrected and corrected signals were evaluated. For each pair of components, spatial similarities were quantified using the absolute Pearson correlation computed across pixels within the shared cortical mask. This resulted in a component-by-component similarity matrix. Optimal one to one matching between components was determined using the Hungarian algorithm to maximize the total spatial correlation across matched pairs.

Spatial overlap between corresponding parcels was also quantified using the Dice similarity coefficient:

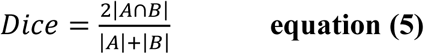

where A and B represent binary masks of matched parcels from the uncorrected and corrected datasets, respectively. Dice scores provide a measure of spatial consistency between parcellation, with higher values indicating greater overlap.

Since spatial correlation and Dice overlap capture complementary aspects of parcel similarity, both metrics were used to classify component stability. Components were classified as stable when both measures indicated strong correspondence (spatial correlation ≥ 0.70 and Dice ≥ 0.50), unstable when either measurement indicated poor correspondence (spatial correlation < 0.50 or Dice < 0.30), and intermediate otherwise. These thresholds were used as operational criteria to separate clearly preserved, clearly altered, and partially preserved functional parcels.

### Functional Connectivity and Network Organization Analysis

Functional connectivity was computed using Pearson correlation between regional fluorescence time series, followed by Fisher z-transformation to enable parametric statistical analysis. Correlation matrices were generated for each recording session from both blue-only and hemodynamic-corrected signals using region-averaged or pixel defined fluorescence traces and summary connectivity metrics were derived from the upper triangle of the z-transformed matrices.

Analyses were performed across cortical regions grouped into functional domains including motor (MOp, MOs), retrosplenial (RSagl, RSPd), somatosensory (SSpll, SSptr, SSpul), and visual (VISa, VISam, VISpm). Pixel based analysis was performed on parcels and segmented by distance. For each animal, connectivity metrics were averaged across sessions to obtain a single representative value per animal.

In addition to pairwise connectivity as a general measurement, graph theoretical analyses were used to further evaluate community structure. Fisher z-transformed FC matrices were converted to weighted graphs by retaining positive edge weights and setting negative weights and diagonal entries to zero. Node strength was defined as:

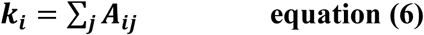

Where *A_ij_* is the positive weighted adjacency matrix. Total graph weight was calculated as:

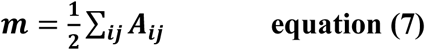

Community structure was estimated using a greedy modularity optimization algorithm. The resulting partition was used to calculate modularity Q. This value should be interpreted as the modularity of the detected greedy partition rather than a guaranteed global maximum. Modularity Q was calculated as:

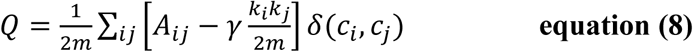

Where *c_i_* and *c_j_* are the community assignments of nodes *i* and *j*. *δ*(*c_i_*, *c_j_*)=1 if nodes are assigned to the same community and 0 otherwise, and *γ* is the resolution parameter. Higher Q values indicate greater within community connectivity than expected under a strength preserving null model.

To complement modularity Q, within community and between community FC were calculated using the detected community assignments. Within community FC was defined as the mean Fisher z-transformed connectivity among node pairs assigned to that same community:

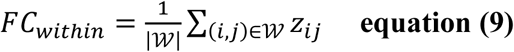

where:

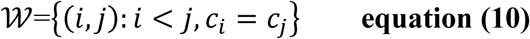

Between community FC was defined as the mean connectivity among node pairs assigned to different communities:

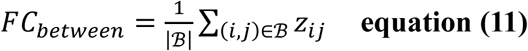

Where:

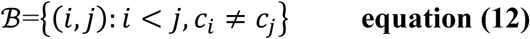

For supplemental topology visualizations, communities detected from the corrected parcel matrices were used as a fixed partition for comparing blue only and hemodynamic corrected networks. Corrected signal community assignments were detected using *γ* = 1.25. Strong edges were defined as parcel-pair Fisher z FC values greater than or equal to the pooled 60^th^ percentile threshold calculated across blue-only and corrected group average matrices for each GCaMP line. Strong edges were classified as within community if both nodes belonged to the same detected community and between communities if nodes belonged to different detected communities. This visualization was used to illustrate whether strong FC edges were broadly distributed across communities or preferentially concentrated within detected communities following hemodynamic correction.

For group-level analyses, connectivity and graph metrics were first calculated for each recording session and then averaged across sessions to obtain one representative value per animal. Blue-only and hemodynamic corrected metrics were compared using paired statistical testing.

### Frequency Domain Analysis

Frequency domain analyses were performed to characterize oscillatory dynamics of widefield calcium signals. Time domain fluorescence traces were transformed into the frequency domain using power spectral density (PSD) estimation based on the Welch method. Spectral analyses were performed within the frequency range of 0.01-9.99 Hz.

For pixel-based analyses, fluorescence signals were extracted from all valid pixels within the cortical mask and reshaped into pixel-wise time series. PSD was computed independently for each pixel and subsequently averaged across pixels within each session. For atlas-based analyses, region averaged fluorescence time series were used and PSD was computed for each region before averaging across regions.

Spectral power was quantified across predefined frequency bands including delta (0.01-4 Hz), theta (4-8 Hz), and lower alpha (8-9.99 Hz). Band specific power was calculated by integrating the PSD within each frequency range:

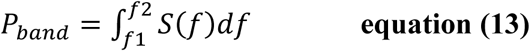

where *S*(*f*) represents the PSD and ⌈*f*_1,_*f*_2_⌉ defines the frequency bounds of each band. The integral was estimated numerically using the trapezoidal rule. Relative power for each band was computed as the proportion of total power within the full frequency range (0.01-9.99 Hz), while absolute power was calculated in linear units and subsequently converted to decibel (dB) scale for reporting.

To quantify the balance between slow and fast oscillatory activity, a fast-to-slow index was defined as:

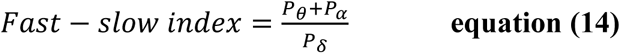

where *P_δ_*, *P_θ_*, *P_α_* correspond the spectral power within the delta, theta, and lower alpha bands, respectively.

All frequency domain metrics were computed separately for blue-only and hemodynamic corrected signals. Session level measurements were first calculated and subsequently averaged across sessions to obtain a single representative value per animal for group level comparisons.

### Multivariate Feature Integration and Principal Component Analysis

Hemodynamic correction was further evaluated within a multivariate framework that integrates spectral, connectivity, and network level metrics into a unified analysis space. Feature vectors were constructed at the session level by combining pixel-based distance-binned functional connectivity across multiple frequency windows, atlas-based connectivity measures, network modularity, and clustering coefficients, relative spectral power (delta, theta, and lower alpha), and fast-slow indices derived from both pixel-based and atlas-based analyses. All features were standardized using z-score normalization prior to dimensionality reduction to ensure comparable scaling across metrics. Missing values were filled with mean imputation.

Principal component analysis (PCA) was applied to the standardized feature matrix to identify dominant axes of variance. Two complementary approaches were used. A combined PCA including all GCaMP lines was performed to place all sessions within a shared feature space, enabling direct comparison across indicators. In parallel, separate PCA models were computed for each GCaMP line to resolve line-specific structure.

The first three principal components were retained for analysis and visualization. Session level coordinates in PCA space were used to compare blue-only and hemodynamic corrected signals. Displacement induced by hemodynamic correction was quantified for each session as Euclidean distance between blue and corrected representation in PC space.

To assess consistency across animals, session level PCA coordinates were averaged within each animal for each signal condition. These animal level representations were used to visualize directional shifts and summarize correction induced changes in GCaMP lines.

### Pupil-brain Coupling Analysis

Relationships between behavioral state and cortical activity were assessed using cross-correlation analysis. Pupil diameter signals were temporally aligned to imaging data and resampled to match the imaging sampling rate (20 Hz). Both pupil and fluorescence signals were preprocessed using Gaussian smoothening (σ = 0.2s), followed by zero-phase Butterworth bandpass filtering (0.02-9.5 Hz) and z-scored.

Cross-correlation functions were computed between pupil signals and regional fluorescence time series for each session. From these functions, multiple metrics were extracted including the zero-lag correlation, the magnitude of and lag of maximum positive correlation peak, and the lag corresponding to the maximum absolute correlation. Peak values and lags were constrained to a ± 5 s window.

Analyses were performed at the regional level, and results were summarized across sessions for each GCaMP line. Mean cross-correlation curves and peak metrics were computed across sessions to assess differences between blue-only and hemodynamic subtracted signals.

### Generalized Linear Modeling of Brain-behavior Relationships

Relationships between behavioral dynamics and cortical activity were quantified using linear modeling approaches. Regional fluorescence signals were modeled as a function of pupil diameter and airball-derived locomotion variables, including forward, lateral, and rotational components.

Pupil diameter was acquired at 40 Hz, whereas imaging data were collected at 20 Hz. Behavioral signals were resampled to the imaging sampling rate using linear interpolation to ensure temporal alignment. Time series were standardized by z-scoring prior to analysis.

Signals were smoothed using a 7 s moving average filter to reduce high-frequency noise and improve model stability. For each region, fluorescence time series were modeled using a Gaussian generalized linear model with an identity link function, equivalent to ordinary least squares regression. The design matrix included pupil diameter and locomotion variables along with an intercept term. Model performance was quantified using the coefficient of determination (R^2^) between predicted and observed fluorescence signals.

Reciprocal relationships were assessed by modeling pupil diameter as a function of cortical activity. Both single-region and multi-region linear models were evaluated, using either individual regional fluorescence signals or multiple regional fluorescence signals as predictor variables. Model performance was again assessed using R^2^, and regression coefficients (β) were extracted to quantify the relative contribution and directionality of individual predictors.

### Functional connectivity, asymmetry, and dynamic network analysis in GBM

Nodal strength was computed from Fisher z-transformed FC matrices as the mean connectivity of each region to all other regions. To quantify tumor related hemispheric effects, connectivity measures were compared between the tumor bearing (ipsilateral) and contralateral hemispheres. In addition to absolute differences, a normalized asymmetry metric was computed as:

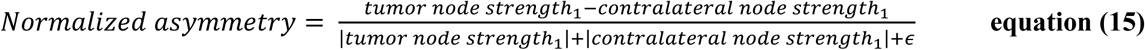

This normalization accounts for differences in signal magnitude and enables comparison of asymmetry across regions and animals.

Dynamic functional connectivity (FC) was computed from region averaged fluorescence time series using a sliding window approach. For each session, hemisphere, and signal type (blue and hemodynamic corrected), pairwise Pearson correlations were calculated within 30 s windows advanced in 2 s steps and transformed using Fisher z. The upper triangle of each FC matrix was vectorized to generate a window level representation of connectivity. These window level FC estimates were used for trajectory analyses and subsequently averaged to obtain session level and animal level measures of region pair connectivity.

To examine the structure of connectivity across regions, window level FC vectors from signals were pooled within each session and hemisphere and underwent principal component analysis (PCA), yielding a shared low-dimensional representation of dynamic FC trajectories. The first three principal components (PC1 and PC2) were used to characterize the primary contributions of connectivity features.

State structure within PCA space was characterized using k-means clustering. Cluster assignments were used to quantify dynamic properties of FC trajectories. For each session and signal type, trajectory length, number of state transitions, number of states visited, and state entropy were computed. Trajectory length was defined as the cumulative Euclidean distance between consecutive windows in principal component space. State entropy was calculated as the Shannon entropy of the distribution of state occupancy:

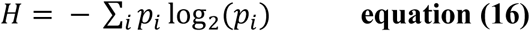

where *p_i_* represents the probability of occupying state *i*. Dwell time metrics were additionally computed to quantify the duration of consecutive occupancy within each state.

To further characterize differences between signals, linear discriminant analysis (LDA) was applied to window level FC vectors. Discriminant weights were sign aligned such that positive values correspond to higher connectivity in corrected signals relative to blue-only signals. Classification was estimated using stratified cross-validation.

All metrics were computed at the session level and averaged across sessions to obtain a single representative value per animal for group level comparisons.

### Deep learning prediction hemodynamic corrected signals (HemoTCN)

A supervised deep learning framework, HemoTCN was developed to predict hemodynamically corrected fluorescence signals directly from blue channel widefield calcium imaging. Training data consisted of dual wavelength recordings acquired from GCaMP6s, GCaMP6s, and jGCaMP8m mice generated by in the Yildirim lab, together with publicly available GCaMP6s recordings from the BraiDynBC dataset^56^. Ground truth corrected signals were generated using the same dual wavelength regression procedure described above for atlas based hemodynamic correction, providing paired blue-only and corrected signals for supervised learning.

To enable cross-laboratory training, recordings from all datasets were harmonized into a unified session level format containing regional fluorescence traces, corrected target signals, anatomical region labels, hemisphere assignments, calcium indicator identity, source laboratory, and sampling frequency. All recordings were resampled to a common temporal resolution (20 Hz) prior to training. Animal level portioning was used to generate training, validation, and test sets, ensuring that recordings from the same animal were never distributed across partitions to prevent data leakage.

The prediction model consisted of a residual temporal convolution network designed to map blue channel fluorescence windows to predicted corrected signals. The architecture contained stacked residual temporal convolutional blocks with exponentially increasing dilation factors to capture both short and long timescale temporal structure within the fluorescence signal. Residual prediction was implemented such that the network estimated a correction term added to the blue input signal, biasing the model toward minimal modification unless hemodynamic contributions were detected. The final model contained approximately 130,000 trainable parameters.

Training was performed using overlapping 256 frame sliding windows extracting from all regions and sessions. To avoid dominance of overrepresented datasets during optimization, balanced minibatch sampling was performed across laboratories and indicator groups. Sliding window segmentation generated 357,500 total windows (263,500 training, 41,250 validation, and 52,750 test windows). Model optimization minimized mean squared error between predicted and ground truth corrected signals using Adam optimizer with early stopping based on validation loss. Full session reconstructions were generated by averaging window predictions across the duration of each recording, producing temporally continuous predicted corrected signals aligned to the original signal.

Model performance was evaluated on held out test sessions using complementary spectral and network level metrics. Spectral recovery was quantified using the fast-slow index (FSI). Functional connectivity recovery was assessed using Fisher z transformed Pearson correlation matrices computed across regions. In addition, the spectral signature of correction was evaluated by comparing log ratio power spectral density curves between blue, true corrected, and model predicted corrected signals. Evaluation metrics were computed at the session level and averaged across animals for group comparisons.

To assess generalization beyond healthy recordings, the trained model was applied without retraining to an independent cohort of jGCaMP8m mice bearing unilateral SB28 glioblastoma allografts. Tumor recordings were processed using the same harmonized session format and inference pipeline as healthy recordings. Predicted corrected signals were subsequently analyzed using the same spectral and network level approaches described above, including hemisphere specific measurements for ipsilateral and contralateral hemispheres.

### HemoTCN Graphical User Interface

To facilitate broad application of deep learning-based correction, we developed HemoTCN, an open-source software platform and graphical user interface (GUI) for prediction of hemodynamic corrected widefield calcium signals from blue-only recordings. The software was implemented in Python as a locally executable Streamlit based application designed for laboratory workstation deployment.

The GUI is organized into multiple analysis modules including: (1) hemodynamic correction inference, (2) correction quality evaluation, (3) transfer learning based fine tuning, (4) de novo model training, and (5) model library management. Native operating system file selection dialogs enable batch loading of imaging sessions and automated dataset ingestion. The software supports multiple neuroscience data formats including MATLAB (.mat), NumPy archive (.npz), and Neurodata Without Borders (.nwb) files through an automated preprocessing pipeline that standardizes recording into an internal representation (**Supplementary Fig. 9**).

Interactive visualization tools were implemented including time domain fluorescence traces, Welch power spectral density overlays, FSI summaries, frequency band power comparisons, and functional connectivity heatmaps. In addition to inference using pretrained models, the interface supports user guided transfer workflows, allowing laboratories with dual wavelength recordings to fine-tune pretrained models or train new models on laboratory specific datasets.

To improve generalization to laboratories not represented in the original training data, the platform additionally incorporates amplitude normalization and optional spectral fine-tuning procedures that adapt model predictions to the statistical properties of new datasets without requiring complete model retraining. We released all the details about HemoTCN in our github page (https://github.com/yyildirimlab/HemoTCN).

## Author Contributions

Conceptualization, T.C. and M.Y.; methodology, T.C., M.E.L, J.H., M.M., K. O., A.S., and M.Y.; software, T.C., and K.O.; validation, T.C., and J.H.; formal analysis, T.C., J.H., and M.E.L.; investigation, T.C., M.E.L., F.B., M.Y.; resources, J.D.L., S.M., and M.Y.; data curation, T.C., J.H., M.E.L.; writing – original draft, T.C. and M.Y.; writing – review & editing, T.C., M.E.L., J.H., M.M., K.O., A.S., J.D.L., S.M., and M.Y.; visualization, T.C., J.H., and M.Y.; supervision, M.Y.; project administration, M.Y.; funding acquisition, M.Y.

## ACKNOWLEDGEMENTS

This work was supported by US National Institute of Health (NIH) grants #R00EB027706 (MY), Case Comprehensive JumpStart Grant #3209 (MY), American Brain Tumor Association Discovery Grant #DG2500080 in memory of Kaitlyn Berg (MY), Cleveland Clinic Research (JDL, MY), Case Western University SOURCE Fellowship (KO), and the Case Comprehensive Cancer Center (JDL)

---

**Supplementary Figure 1.**
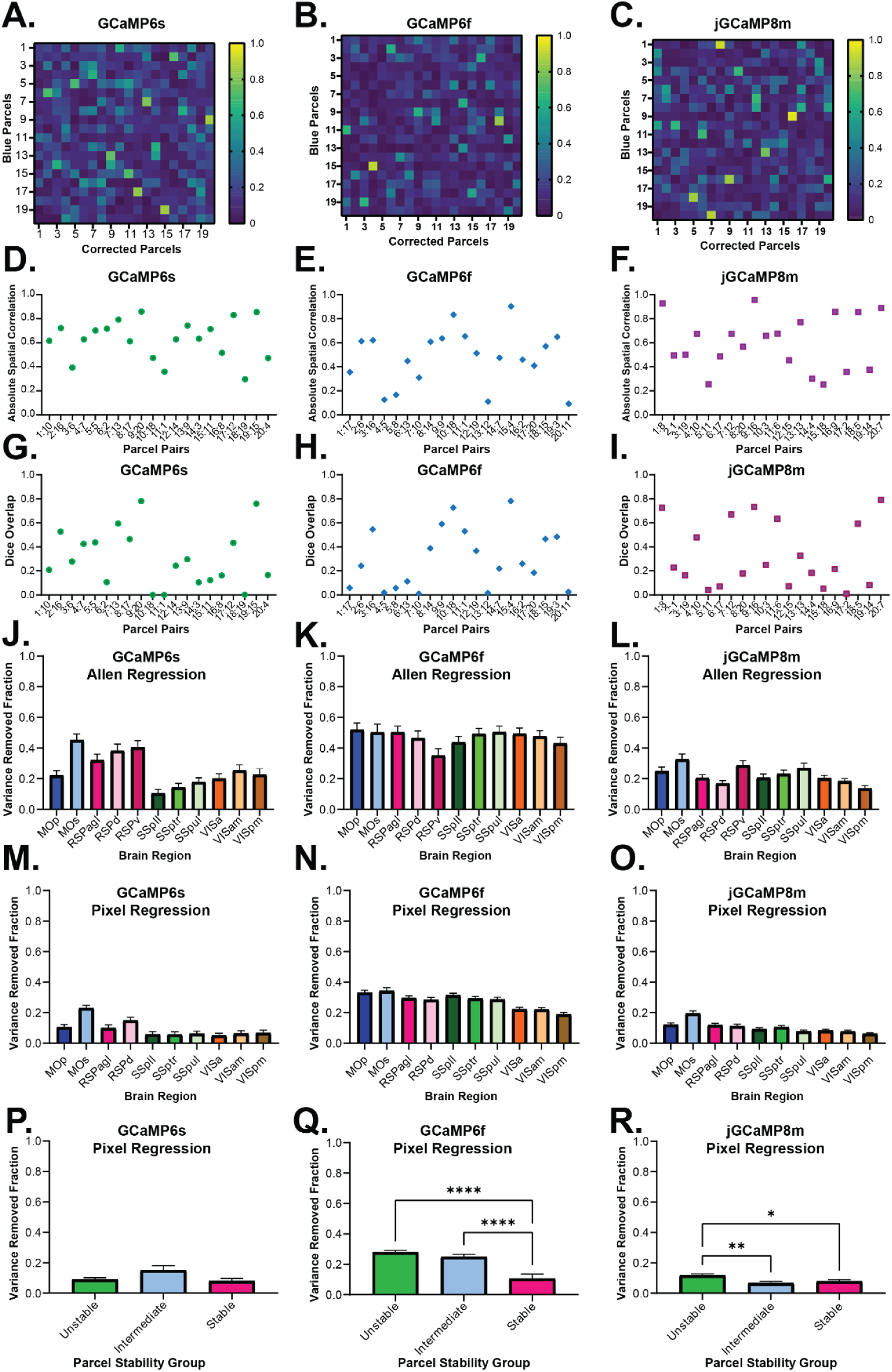
Functional parcellation similarity and variance removed fraction across GCaMP lines. **(A-C)** Heatmaps showing pairwise spatial similarity between blue-derived and hemodynamic corrected ICA parcels for GCaMP6s, GCaMP6f, and jGCaMP8m mice. **(D-F)** Absolute spatial correlation between matched parcel pairs for each GCaMP line. **(G-I)** Dice overlap quantifying spatial agreement between matched parcels. **(J-L)** Variance removed fraction (VRF; mean ± SEM) across cortical regions following atlas-based regression. **(M-O)** VRF (mean ± SEM) for the same regions following pixel-based regression. **(P-R)** VRF (mean ± SEM) stratified by parcel stability group (unstable, intermediate, stable) for pixel-based regression. Statistical significance was assessed using ordinary one-way ANOVA followed by Tukey’s multiple comparisons test. Significance is indicated as *p* < 0.05 (*), *p* < 0.01 (**), *p* < 0.001 (***), and *p* < 0.0001 (****).

**Supplementary Figure 2.**
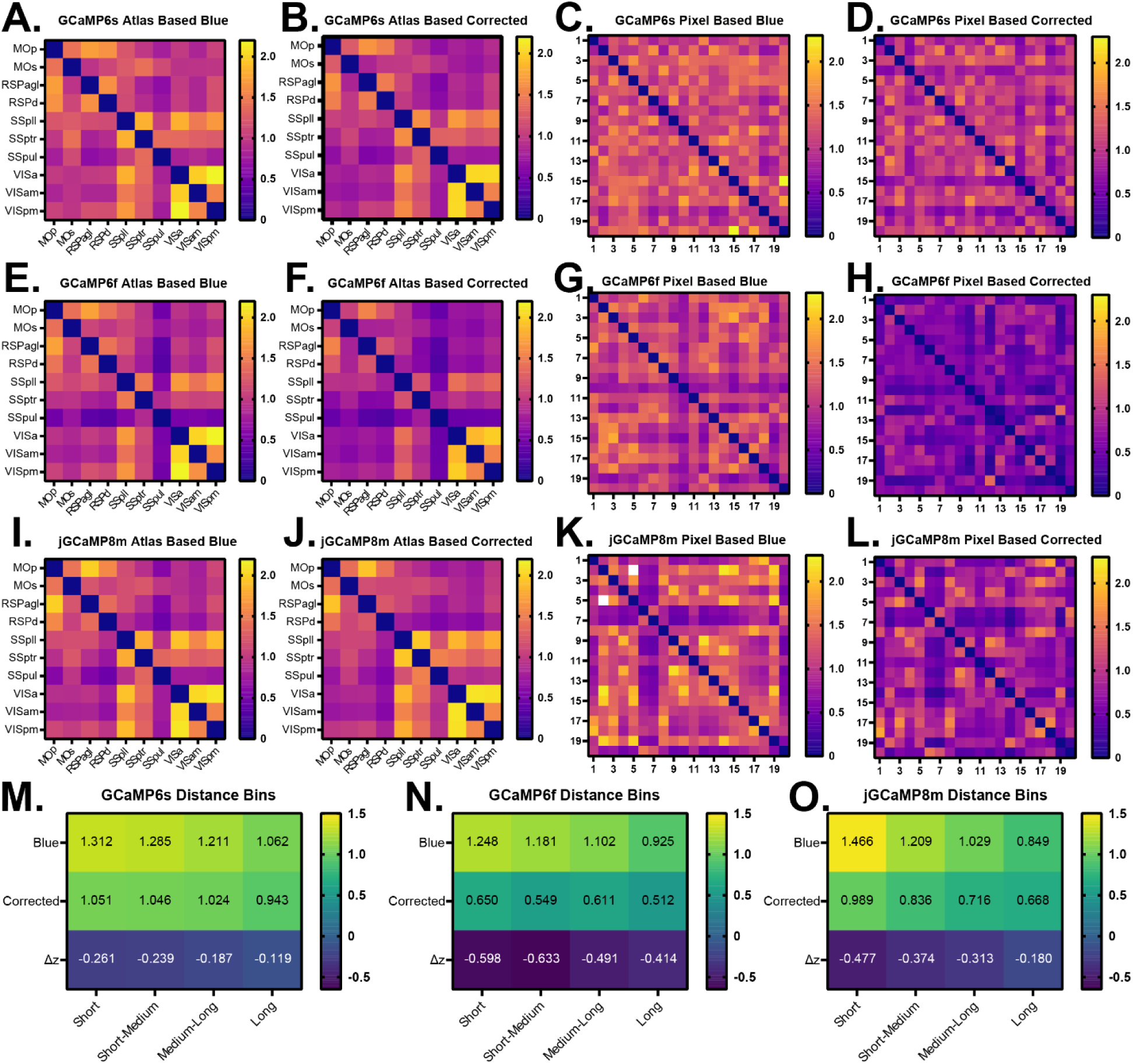
Functional connectivity matrices and distance-resolved changes across GCaMP lines. **(A-D)** Functional connectivity matrices (Fisher z) for GCaMP6s showing **(A)** Atlas based blue, **(B)** Atlas-based corrected, **(C)** Pixel-based blue and **(D)** Pixel-based corrected signals. **(E-H)** Functional connectivity matrices for GCaMP6f showing atlas-based **(E)** blue, **(F)** corrected and pixel-based **(G)** blue and **(H)** corrected signals. **(I-L)** Functional connectivity matrices for jGCaMP8m showing atlas-based **(I)** blue, **(J)** corrected and pixel-based **(K)** blue and **(L)** corrected signals. **(M-O)** Distance resolved connectivity summaries (mean ± SEM) for short short-to-medium medium-to-long and long-range connections for **(M)** GCaMP6s, **(N)** GCaMP6f, and **(O)** jGCaMP8m including blue-only corrected and Δ connectivity (corrected-blue).

**Supplementary Figure 3.**
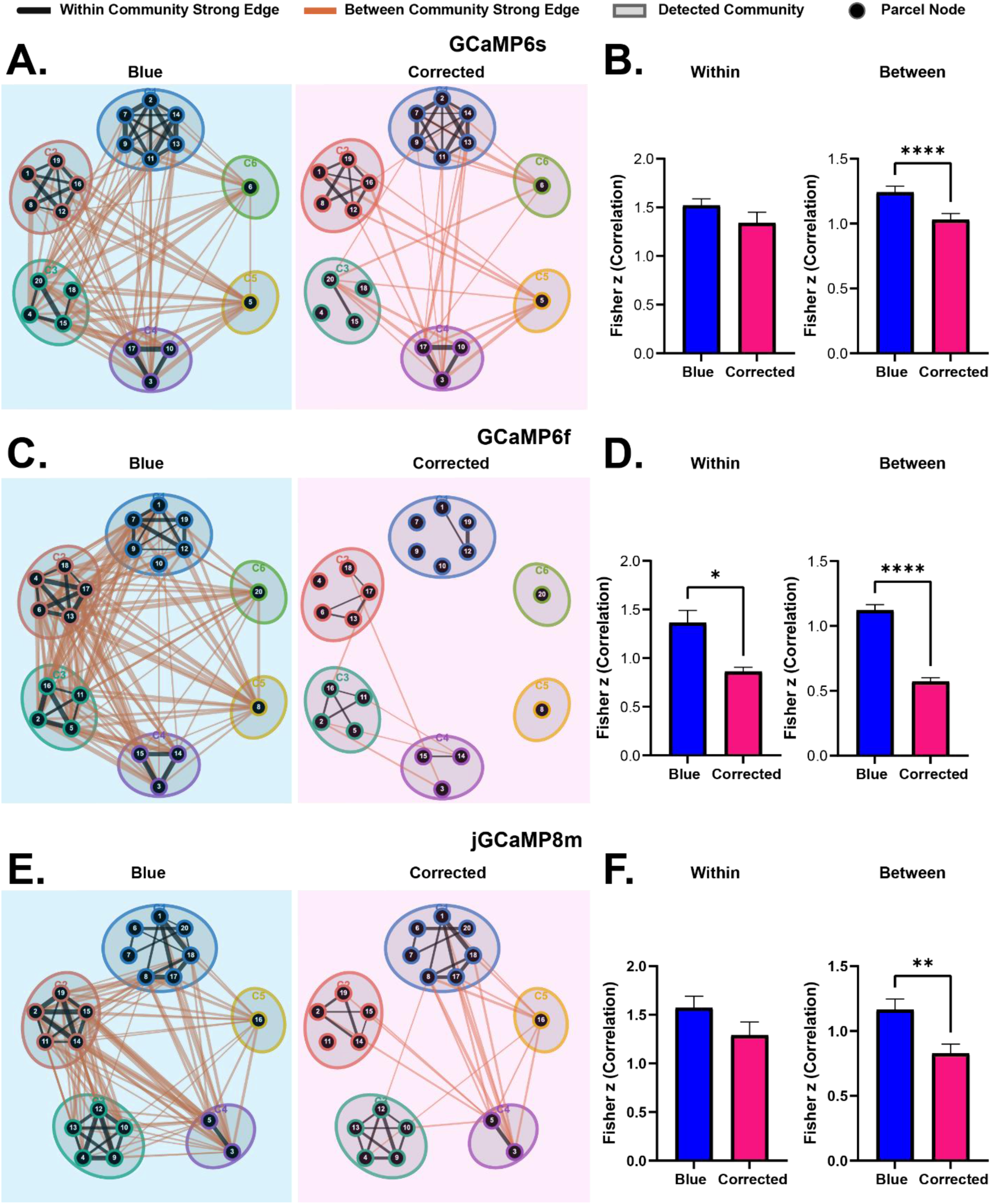
Hemodynamic correction reduces cross-community connectivity and increases modular segregation in pixel-derived parcel networks. Clustered topology plots for GCaMP lines comparing blue (left) and hemodynamic corrected (right) parcel functional connectivity matrices. Nodes represent pixel derived parcels, gray regions indicate detected community connections, and orange edges indicate strong between community connections. Strong edges were defined as parcel-pair Fisher z FC values greater than or equal to the pooled 60^th^ percentile threshold calculated across blue and corrected group average matrices for each line. **(A)** Topology plots for GCaMP6s. Strong within/between count was 30/69 for blue and 26/27 for corrected. **(B)** GCaMP6s quantification of mean within community connectivity (left) and between community connectivity (right). **(C)** Topology plots for GCaMP6f. Strong within/between count was 27/109 for blue vs. 11/5 for corrected. **(D)** GCaMP6f quantification of within (left) and between (right) community connectivity. **(E)** Topology plots for jGCaMP8m. Strong within/between count was 33/77 for blue and 24/18 for corrected. **(F)** jGCaMP8m quantification of within (left) and between (right) community connectivity. Bars indicate mean ± SEM. Significance was assessed using paired two tailed t-test. Significance is indicated as *p* < 0.05 (*), *p* < 0.01 (**), *p* < 0.001 (***), and *p* < 0.0001 (****).

**Supplementary Figure 4.**
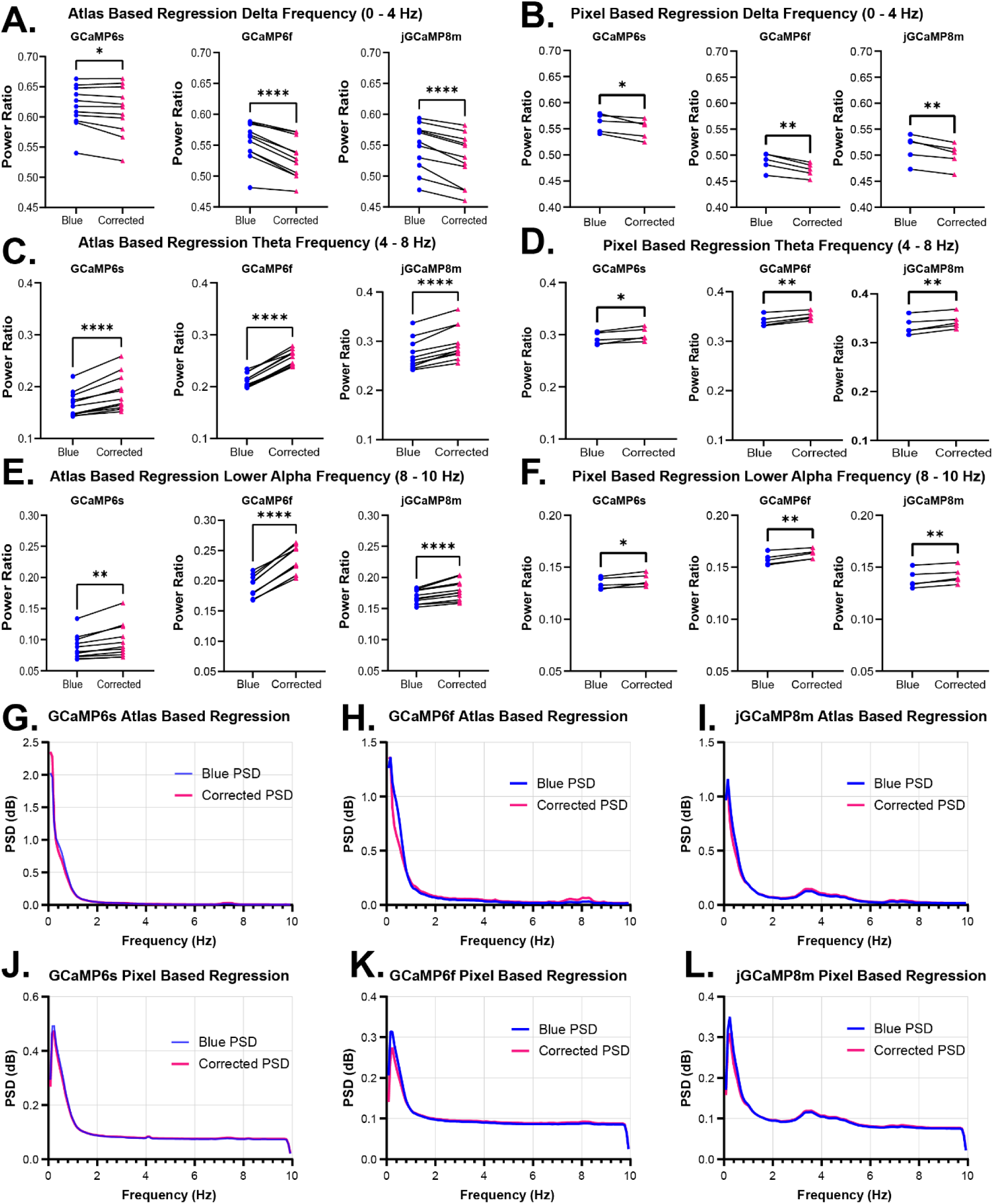
Frequency band specific power changes and power spectral density following hemodynamic correction. **(A-B)** Delta band relative power for **(A)** atlas-based and **(B)** pixel-based analyses across (left) GCaMP6s, (middle) GCaMP6f and (right) jGCaMP8m. Hemodynamic correction significantly reduced delta band power across all lines (paired two-tailed t-test; atlas: p = 0.0359, <0.0001; pixel: p = 0.0261, 0.0018, 0.0051 for 6s, 6f, and 8m respectively). **(C-D)** Theta band relative power for **(C)** atlas-based and **(D)** pixel-based analyses, showing significant increases following correction across GCaMP lines (all p ≤ 0.0243). **(E-F)** Lower alpha band relative power for **(E)** atlas-based and **(F)** pixel-based, also showing significant increases with correction (all p ≤ 0.0300). For atlas-based, each point represents a brain region averaged across five animals and six sessions; for pixel based, each point represents an individual animal averaged across six sessions. Bars indicate mean ± SEM. **(G-I)** Power spectrum density (PSD) curves for atlas based across **(G)** GCaMP6s, **(H)** GCaMP6f, and **(I)** jGCaMP8m, showing reductions in low frequency power and relative increases in higher frequency components with correction. **(J-L)** PSD curves for pixel-based analyses across **(J)** GCaMP6s, **(K)** GCaMP6f, and **(L)** jGCaMP8m, demonstrating similar but more pronounced spectral shifts compared to atlas based. Significance is indicated as *p* < 0.05 (*), *p* < 0.01 (**), *p* < 0.001 (***), and *p* < 0.0001 (****).

**Supplementary Figure 5.**
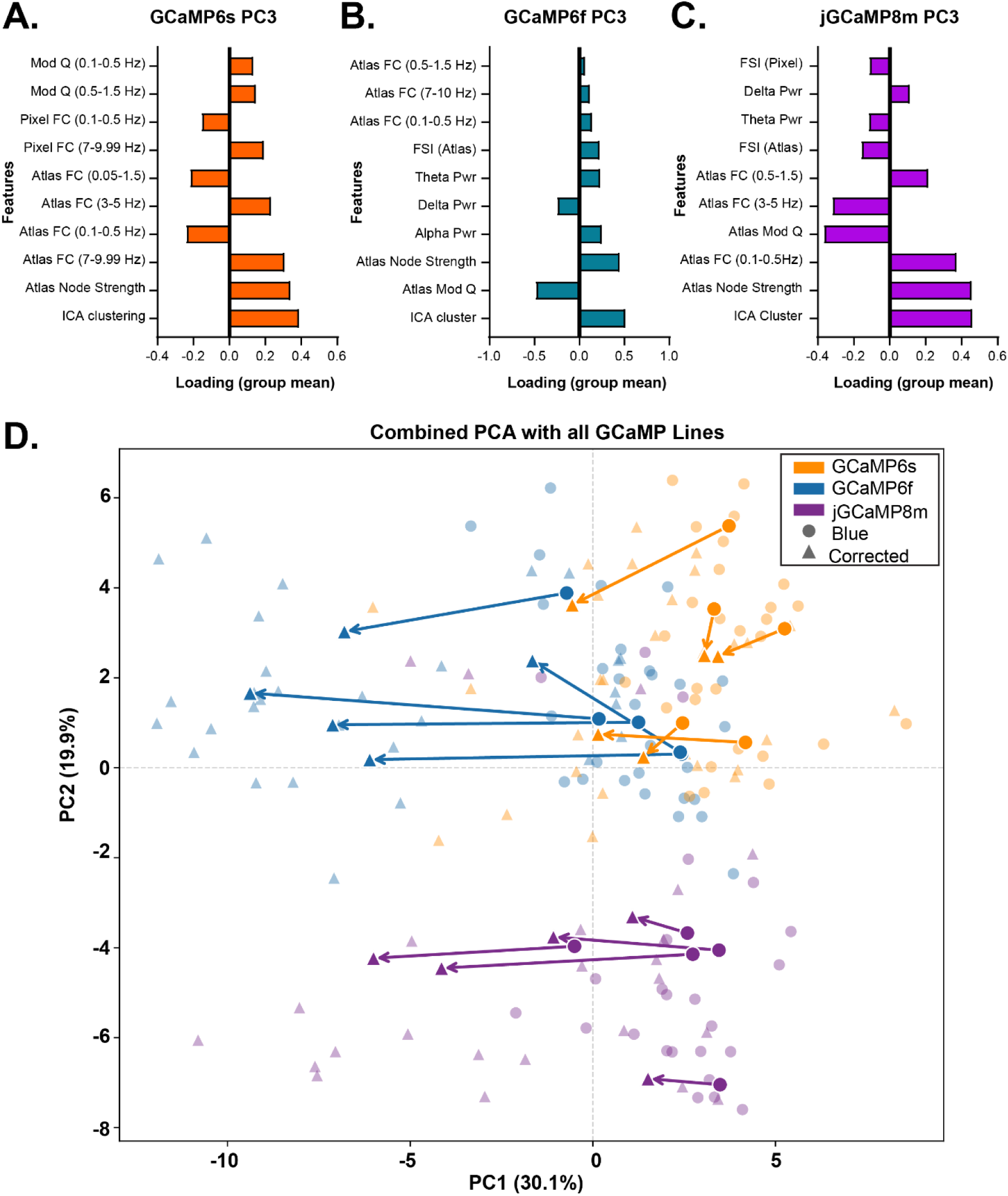
Principal component structure and cross-line PCA comparison. **(A-C)** Feature loadings for the third principal component (PC3) for **(A)** GCaMP6s, **(B)** GCaMP6f, and **(C)** jGCaMP8m, highlighting line specific contributions of network and frequency features to higher order variance structure. **(D)** Combined PCA across all GCaMP lines. Semi-transparent points represent individual recording sessions, while solid markers indicate animal-averaged values. Circles denote blue only signals and triangle denote hemodynamic corrected signals. Arrows indicate within animal transitions from blue only to corrected states in PCA space. The first two principal components explain 30.1% (PC1) and 19.2% (PC2) of the total variance.

**Supplementary Figure 6.**
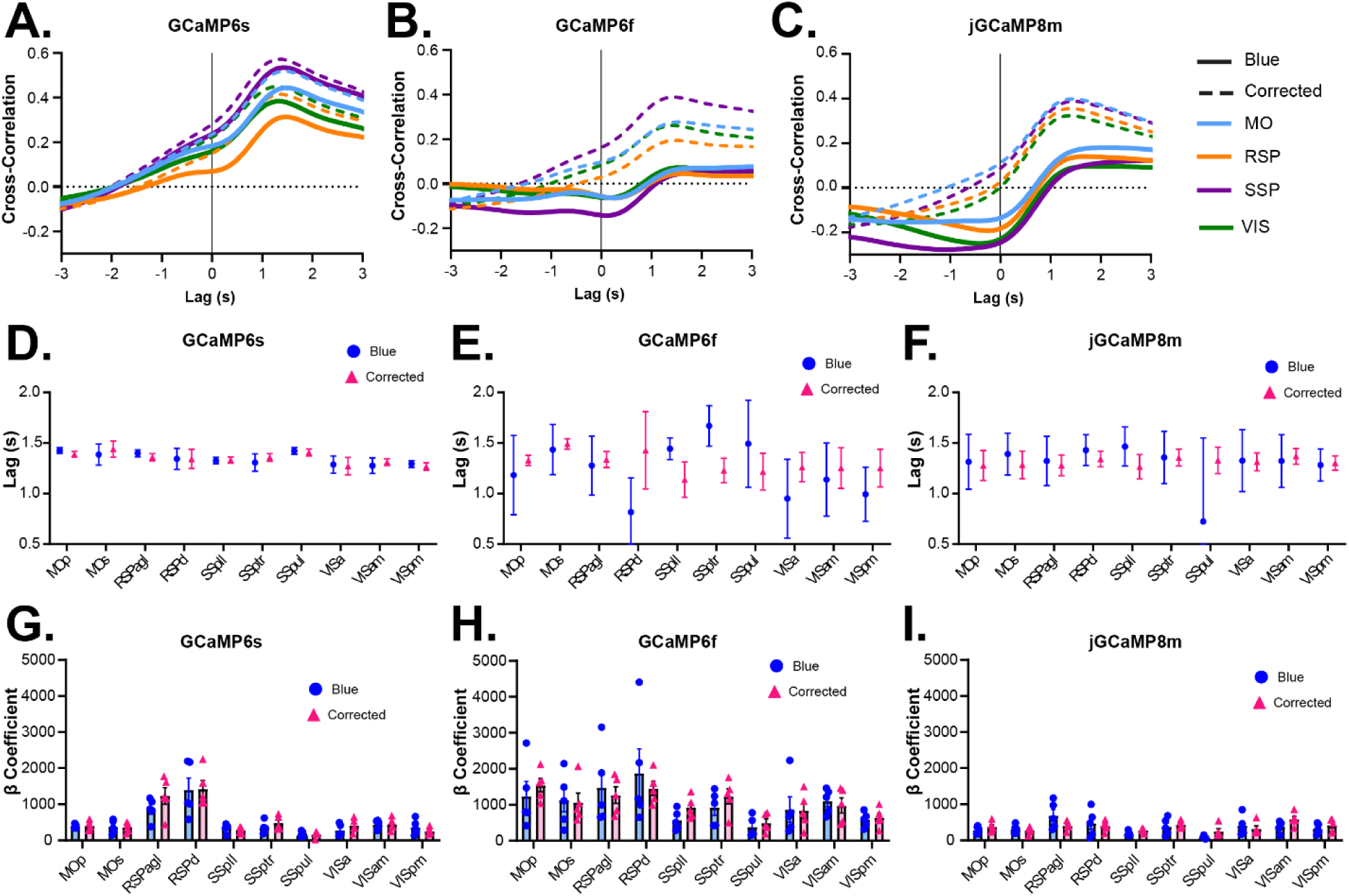
Pupil-brain coupling and temporal relationships across GCaMP lines. Mean cross-correlation functions between pupil diameter and regional fluorescence signals for **(A)** GCaMP6s, **(B)** GCaMP6f, and **(C)** jGCaMP8m. Solid lines represent blue-only signals and dashed lines represent hemodynamic-corrected signals. Cross-correlations are shown as the function of lag (seconds), with positive lags indicating pupil-leading relationships. **(D-F)** Lag at peak cross-correlation across cortical regions for **(D)** GCaMP6s, **(E)** GCaMP6f, and **(F)** jGCaMP8m, highlighting region specific differences in the temporal relationship between pupil dynamics and cortical activity. **(G-I)** Regression coefficients (β) from linear models predicting pupil diameter from regional fluorescence signals for **(G)** GCaMP6s, **(H)** GCaMP6f, and **(I)** jGCaMP8m, indicating the contribution of individual brain regions to pupil dynamics. All values are averaged across recording sessions; points represent individual animals and bars indicate mean ±SEM.

**Supplementary Figure 7.**
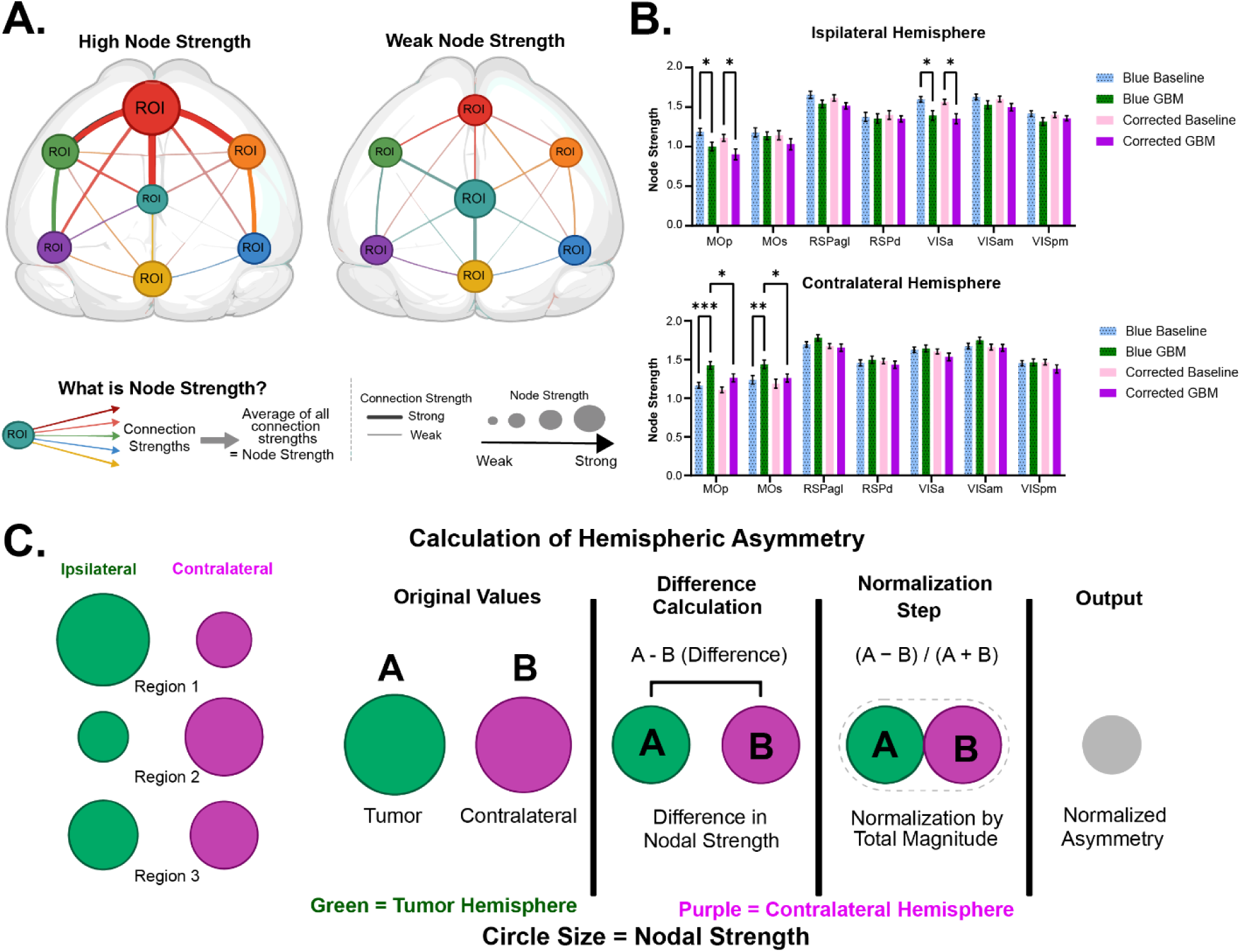
Node strength and hemisphere asymmetry in GBM. **(A)** Schematic illustrating nodal strength, defined as the mean functional connectivity of a region to all other regions and conceptual differences between high and low connectivity nodes. **(B)** Regional nodal strength across baseline and GBM conditions or blue-only and hemodynamic-corrected signals in the ipsilateral (top) and contralateral (bottom) hemispheres. Hemodynamic correction altered nodal strength in a region and hemisphere dependent manner, with significant effects observed in motor and visual regions in the ipsilateral hemisphere and more pronounced reductions in the contralateral hemisphere (two-way ANOVA with Tukey’s multiple comparisons test). **(C)** Schematic illustrating the calculation of hemispheric asymmetry, including absolute and normalized asymmetry metrics. The ipsilateral is shown in green and contralateral hemisphere in purple.

**Supplementary Figure 8.**
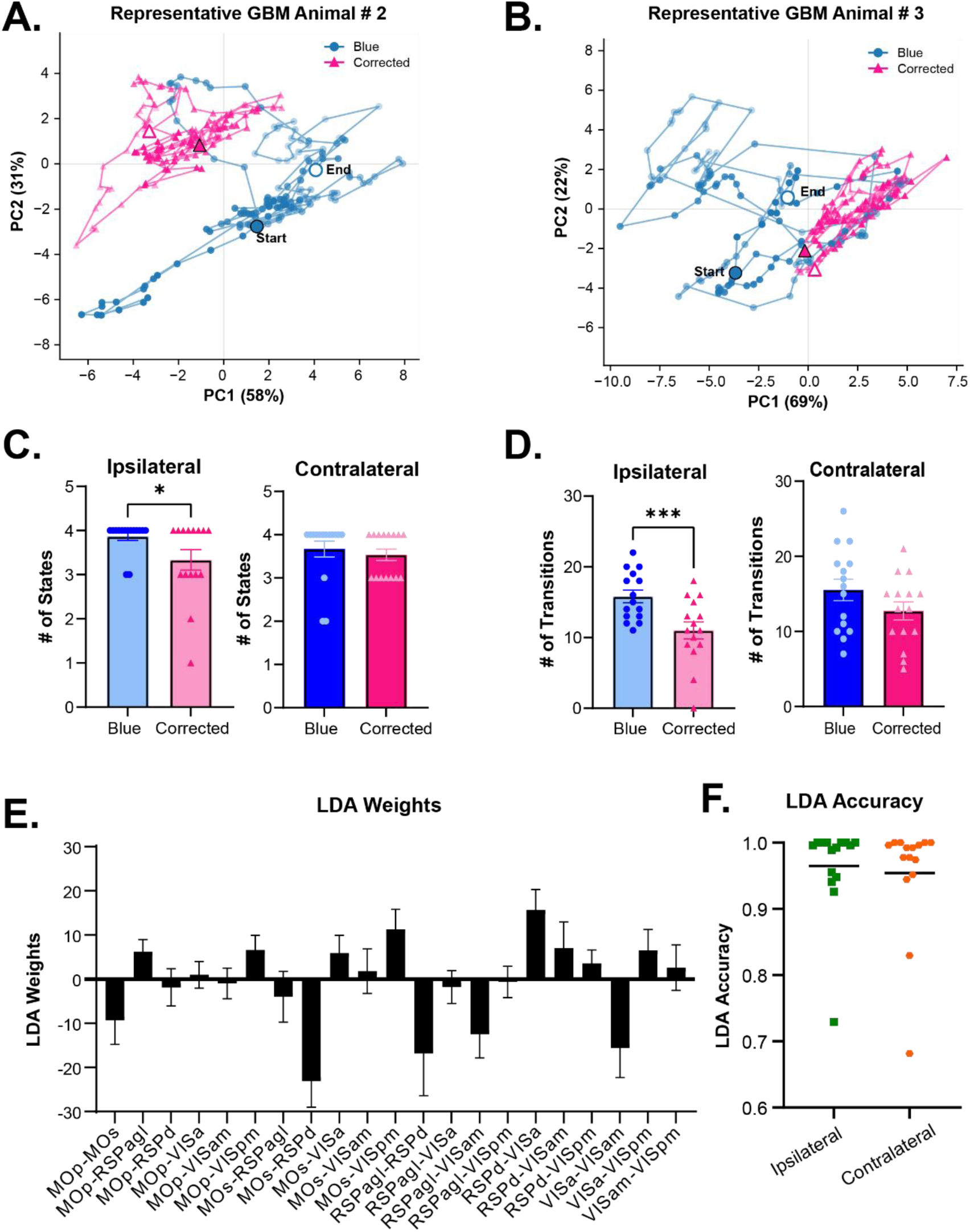
Additional characterization of dynamic functional connectivity states in GBM. **(A-B)** Representative PCA trajectories for two additional animals, showing blue-only (blue) and hemodynamic-corrected (pink) signals. The first two principal components explain 58% and 31% variance in A and 69% and 22% in B. **(C)** Number of k-means states calculated in (left) ipsilateral (p = 0.0266) and (right) contralateral hemispheres (p = 0.4985). **(D)** Number of state transitions between consecutive windows in (left) ipsilateral (p=0.0006) and (right) contralateral hemispheres (p = 0.0632) **(E)** Linear discriminant analysis (LDA) weights identifying region pair contributions that distinguish blue-only and corrected functional connectivity states. **(F)** LDA classification accuracy for distinguishing blue-only and corrected states, demonstrating separability of signals in dynamic functional connectivity space. Points represent individual animals; bars indicate mean ± SEM. Statistical analysis utilized paired two-tailed t-test. Significance is indicated as *p* < 0.05 (*), *p* < 0.01 (**), *p* < 0.001 (***), and *p* < 0.0001 (****).

**Supplementary Figure 9.**
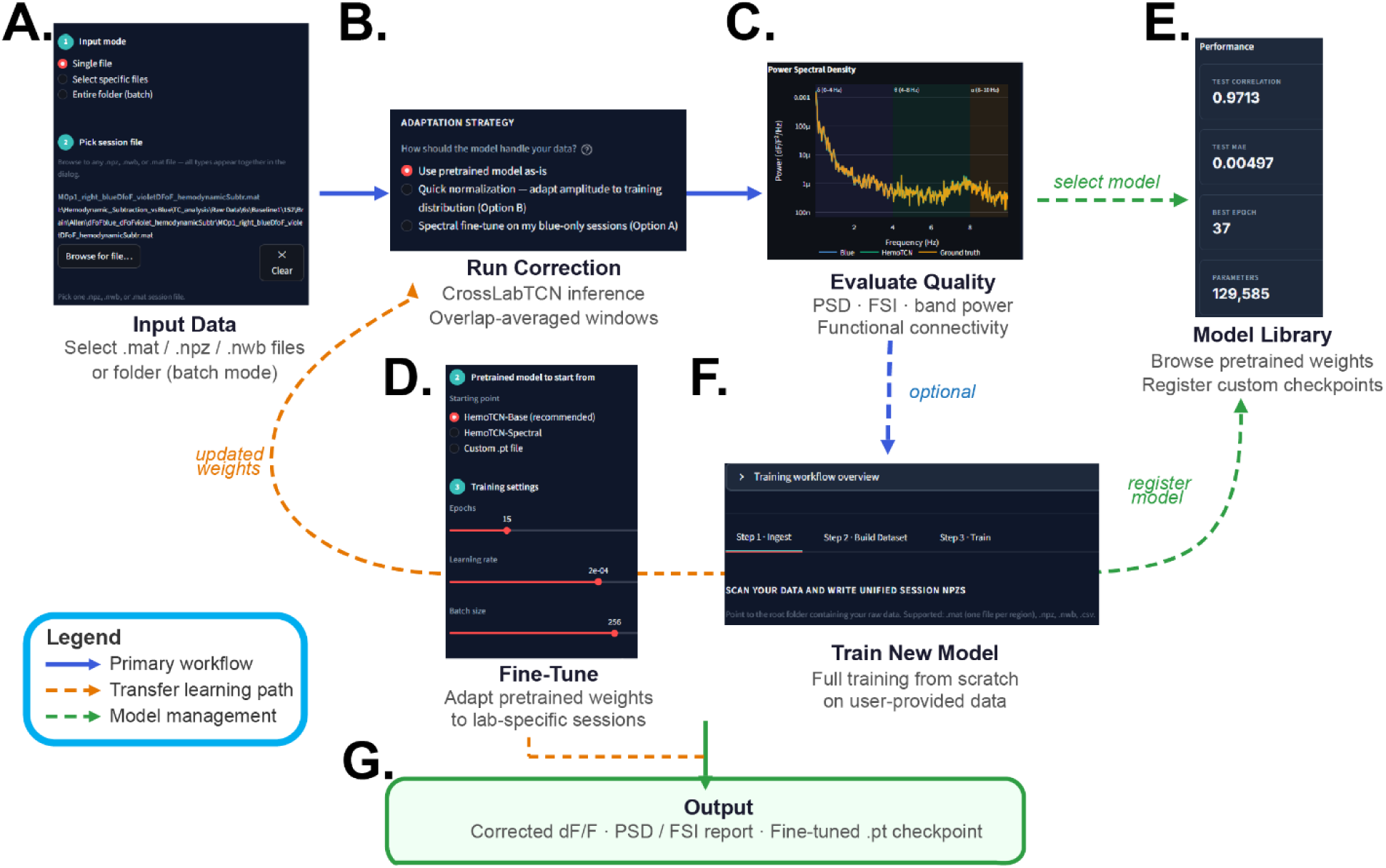
HemoTCN graphical user interface and deep learning workflow for hemodynamic correction. **(A)** Input data selection interface. Users specify imaging sessions as individual files (.mat, .npz, or .nwb) or as folders for recursive batch processing using native operating system file dialogs. **(B)** Hemodynamic correction interface workflow. The multi-lab model processes regional blue channel fluorescence signals using overlapping 256 frame sliding windows with reconstruction. Users specify metadata including anatomical region, hemisphere, calcium indicator, and laboratory identity prior to inference. **(C)** Correction quality evaluation interface displaying interactive visualization including fluorescence traces, PSD estimates, FSI, band power comparisons, and functional connectivity before and after correction. **(D)** Transfer learning and fine-tuning interface enabling adaption of pretrained model weights to laboratory specific dual wavelength recordings. Updated checkpoints are subsequently available for inference. **(E)** Model library interface showing pretrained and user guided checkpoints together with architecture metadata, model performance metrics, and installation status. **(F)** Train from scratch workflow enabling full model training on user provided datasets. Resulting checkpoints can subsequently be registered within the model library for future inference workflows. Blue arrows indicate the primary inference workflow, orange arrows indicate transfer learning workflows, and green arrows indicate model management operations. **(G)** Output is corrected df/f, PSD, and fine-tuned checkpoints that can be downloaded.

## Notes

https://github.com/yyildirimlab/HemoTCN

### Competing Interest Statement

The authors have declared no competing interest.

